# H1.3 depletion in AML cells prompts H1.2 redistribution, chromatin remodeling and cell cycle defects

**DOI:** 10.1101/2025.07.28.667197

**Authors:** Clara Tellez-Quijorna, Lia N’Guyen, Núria Serna-Pujol, Maeva Rodies, Julien Vernerey, Albert Jordan, Estelle Duprez

## Abstract

Linker histone H1 variants play critical, yet distinct, roles in chromatin organization and gene regulation. However, very little is known about their specificity in cancer cells. In this study, we investigated the specificity of the H1.3 variant in acute myeloid leukemia (AML) cells. Through chromatin mapping and transcriptomic analyses, we revealed that H1.3 was enriched explicitly in regions with a high GC content and co-localized with the repressive mark H3K27me3, supporting its role in chromatin compaction and transcriptional repression. Knockout of H1.3 induced specific changes in gene expression profiles and chromatin dynamics, characterized by the relocalization of H1.2, which was redistributed from its usual chromatin regions to the freed H1.3 regions. Consequently, locus-specific chromatin alterations associated with interferon-related signalling pathways and cell cycle deregulation were observed. Our findings emphasized an important and locus-specific function of H1.3 in AML cells. Overall, our study revealed a mechanistic connection between H1 variant imbalance, immune response activation, and cell cycle regulation, with implications for our understanding of epigenetic regulation in cancer cells.

## INTRODUCTION

Chromatin is a fundamental macronucleoprotein complex that organizes the eukaryotic genome, controlling the accessibility of genomic information and some crucial biological processes, including DNA replication, transcription, the cell cycle, and DNA repair (Chen et al., 2024; Dai et al., 2020; Kornberg, 1974). Thus, chromatin dynamics are primarily influenced by epigenetic modifications, which alter its structure and function (Felsenfeld, 1978; Reid et al., 2017; Wanner & Formanek, 2000). The basic repeating unit of chromatin is the nucleosome, which contains 145-147 base pairs (bp) of DNA wrapped in a left-handed 1.65 turns around an octameric histone complex. This octamer is formed by two copies of the core histones H2A, H2B, H3, and H4, organized as an (H3-H4)_2_ tetramer flanked by two (H2A-H2B) dimers (Arents & Moudrianakis, 1995; Olins & Olins, 1974; Oudet et al., 1975). Nucleosomes are interconnected by short DNA segments (20-90 bp) and the linker histone H1, which binds to the entry-exit sites of the nucleosome and plays an essential role in chromatin compaction and higher-order folding (Fyodorov et al., 2018; Noll & Kornberg, 1977; Simpson, 1978).

The linker histone H1 constitutes the most diverse and heterogeneous histone family compared with the highly conserved core histone proteins. As core histones, linker histone H1 can be further divided into different variants, dependent on the organism, in a dependent or independent-replicative manner. In humans, there are seven somatic H1 variants, from H1.1 to H1.5, H1.0, and H1.10. The replication-dependent variants (H1.1–H1.5) are predominantly expressed during the S-phase of the cell cycle, while H1.0 and H1.10 are expressed independently of replication. Among these variants, H1.2 to H1.5 are predominantly present in all tissues, whereas H1.1 displays tissue-specific expression. The independent somatic variants are mainly observed in differentiated cells (Happel & Doenecke, 2009; Prendergast & Reinberg, 2021; Talbert et al., 2012). The classical concept of the linker histone H1 has revolved around its role in stabilizing nucleosomes and promoting the condensation of higher-order chromatin structures (Fyodorov et al., 2018). As this histone family exhibits greater sequence variability across species and variants than the well-conserved core histones, a key challenge is to determine whether this variability confers chromatin specificity to the variants. For many years, the lack of specific antibodies for chromatin immunoprecipitation sequencing (ChIP-seq) has limited our ability to explore the role of H1 variants in chromatin dynamics fully. Nevertheless, several studies have demonstrated that different H1 variants perform specific functions in both chromatin structure and the regulation of various cellular processes (Behrends & Engmann, 2020; Fernández-Justel et al., 2022; Izquierdo-Bouldstridge et al., 2017; Izzo et al., 2013; Lai et al., 2021; Salinas-Pena et al., 2024a; Sancho et al., 2008), although our understanding remains limited.

In addition to their structural and regulatory functions, linker histones have been connected to various diseases (Duce et al., 2006; Tremblay et al., 2022; Ye et al., 2017), including cancer (Bauden et al., 2017; Bonner et al., 2023; Chen et al., 2023; Khachaturov et al., 2014), highlighting their biological importance. In this context, we previously described a novel epigenetic biomarker, H3K27me3*HIST1*, in patients with acute myeloid leukemia (AML) (Tiberi et al., 2015). This epigenetic biomarker is characterized by a focal enrichment of the repressive histone mark H3K27me3 within the major histone gene cluster (the *HIST1* cluster). This epigenetic alteration affected 11 histone genes in the *HIST1* cluster, including the single-copy linker histone H1.3. Consequently, AML patients who were H3K27me3*HIST1*-positive exhibited lower H1.3 expression and a favorable outcome (Garciaz et al., 2019). These findings suggested an unexplored role for the H1.3 linker histone in AML progression.

In this study, we investigated the functional role of H1.3 in AML cells using a CRISPR-Cas9-mediated knockout (KO) model in the *NPM1*-mutated OCI-AML3 cell line. Our findings reveal that the loss of this linker histone variant leads to a reorganization of transcriptional programs, including downregulation of cell cycle genes and upregulation of interferon-stimulated genes, as well as local changes in chromatin accessibility. Our results indicate that upon H1.3 depletion, the linker histone variant H1.2 partially redistributes across the genome. Our work demonstrates that the absence of H1.3 triggers chromatin and transcriptional changes, highlighting the non-redundant role of H1 variants in leukemia biology and may account for the favorable outcome of H3K27me3*HIST1*-positive patients.

## RESULTS

### H1.3 histone variant depletion leads to an upregulation of H1.0 histone variant in leukemic cells

To study the role of H1.3 in AML, we chose to generate a CRISPR-Cas9 KO of H1.3 in the *NPM1*-mutated AML cell line OCI-AML3 (**Fig. 1A**), as H1.3 deregulation was observed in *NPM1* mutant AML patients (Garciaz et al., 2019). We generated H1.3 KO clones, which were analyzed by RT-qPCR and western blot (**Fig. 1A**). We then identified the genetic alteration leading to the absence of H1.3 expression in these clones by DNA sequencing (**Appendix Fig. S1A**). Despite the diversity of H1s, several studies have shown that eliminating a single variant did not alter overall levels of total H1 due to compensatory effects by the other H1 variants (Drabent et al., 2000; Fan et al., 2001, 2003). To determine whether the absence of H1.3 caused a change in the abundance of other H1 proteins, we studied the level of other H1 variants under H1.3 KO. Our RT-qPCR and western blot analyses showed that of the replication-dependent H1 variants, H1.2, H1.4, and H1.5 (**Fig. 1B**, **C**) were expressed in our AML cell lines. However, *H1.4* mRNA was less expressed, while *H1.1* mRNA (**Appendix Fig. S1B**) was barely detectable, suggesting differences in levels between the different linkers H1. Nevertheless, deletion of H1.3 did not alter the protein levels of the replication-dependent H1 variants H1.2, H1.4, and H1.5 (**Fig. 1B**, **C**). Interestingly, we observed an upregulation of the replication-independent linker variant H1.0 at both the mRNA and protein levels (**Fig. 1B**, **C**). Similar upregulation has been reported in breast cancer cell lines following knockdown of H1.2, H1.4, or multiple H1 variants (multi-H1KD) (Izquierdo-Bouldstridge et al., 2017; Sancho et al., 2008). These results suggest that the absence of H1.3 in the OCI-AML3 cells context may be compensated for by the upregulation of the cell-cycle independent H1.0 variant.

**Figure 1.**
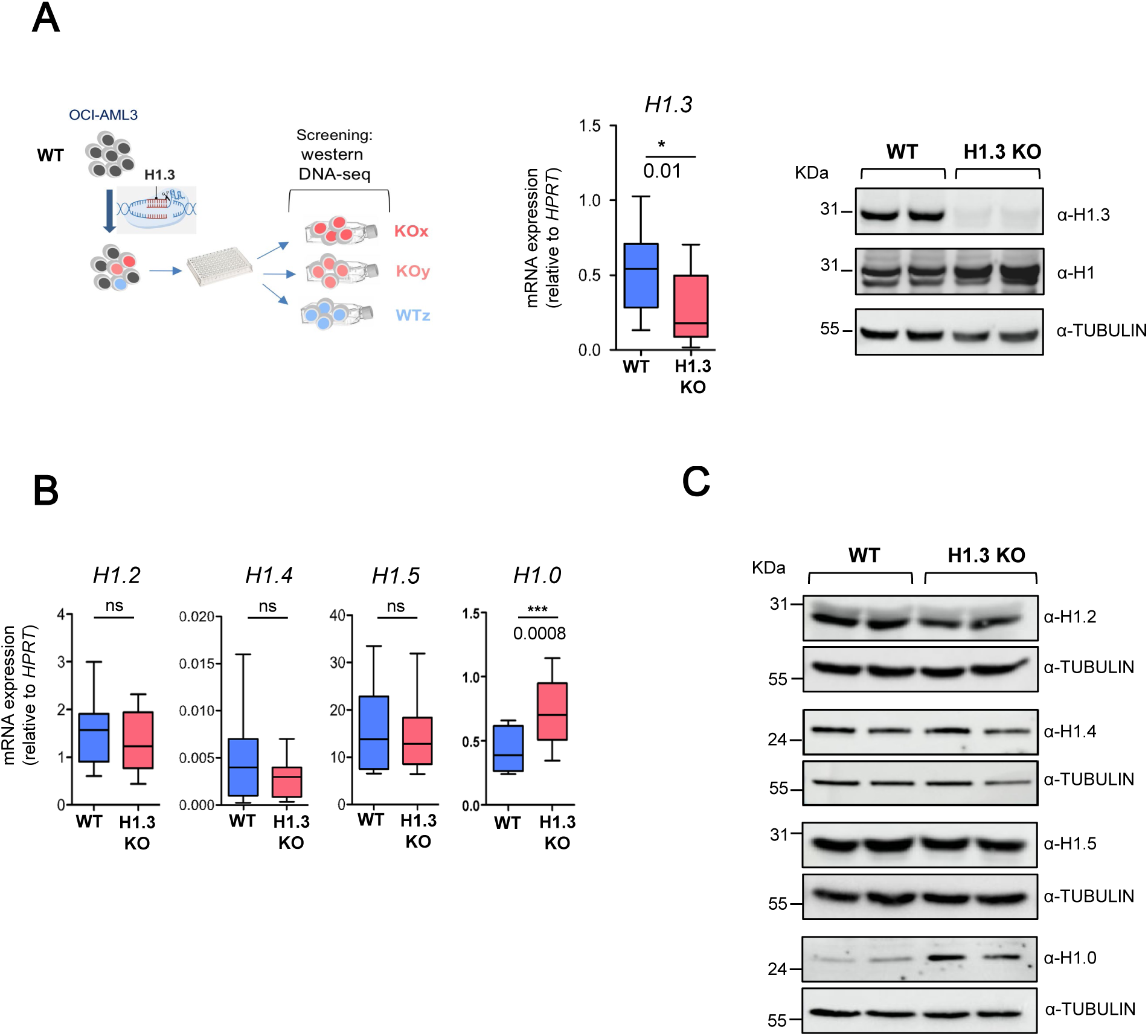
H1.3 knockout validation and analysis of linker histone variant expression in OCI-AML3 cells. (**A**) Characterization of CRISPR-Cas9 H1.3 OCI-AML3 KO clones. Upper, the scheme of clone selection. In the lower left panel, H1.3 mRNA levels were quantified by RT-qPCR and normalized to the housekeeping gene *HPRT*. In the lower right panel, a western blot showing the absence of *H1.3* protein expression in KO clones. (**B-C**) Histone variants H1.2, H1.4, H1.5 and H1.0 analysis upon H1.3 KO. In **B**, *H1.2*, *H1.4*, *H1.5 and H1.0* histone variants were quantified by RT-qPCR in 2 WT and 2 H1.3 KO OCI-AML3 cell lines. mRNA levels were normalized to *HPRT*. Data represent mean ± s.e.m. of n = 6 biological replicates. Statistical significance was estimated using an unpaired two-tailed t-test * p < 0.05; **p < 0.005; *** p < 0.001; ns = no significance. In **C**, H1.2, H1.4, H1.5 and H1.0 histone variants were quantified by western blot in 2 OCI-AML3 WT and 2 H1.3 KO cell lines. TUBULIN was used as a loading control.

### H1.3 depletion modifies transcriptional programs and chromatin accessibility in AML cells

As a linker histone, H1.3 is essential for chromatin compaction and regulation of DNA accessibility (Healton et al., 2019; Prendergast & Reinberg, 2021; Liu et al., 2023), so we investigated the impact of its absence on chromatin accessibility by ATAC-sequencing in both the OCI-AML3 WT and H1.3 KO clones. The genome-wide analysis did not reveal significant alterations in overall chromatin accessibility across the genome (**Appendix Fig. S2A**). However, differential analysis revealed regions modulated in their chromatin accessibility upon H1.3 depletion. A gain in chromatin accessibility was observed in 214 regions in the H1.3 KO condition, while 78 regions were more accessible in the WT condition, suggesting a looser chromatin state in the WT (**Appendix Fig. S2B**; **Table S4**).

Next, to explore the H1.3 depletion effect on gene expression, we performed RNA sequencing in WT and H1.3 KO conditions. We identified 1,139 differentially expressed genes (DEGs; at least Padj <=0.05, absolute (FC) > 1.5), of which 770 genes were upregulated in the KO and 369 genes in the WT condition (**Fig. 2A**; **Table S5**). Upregulated genes in the KO condition exhibited enriched gene signatures related to defense against viruses and type I interferon signaling, indicating a specific response induced by H1.3 depletion. Conversely, the upregulated genes in the WT condition showed enrichment in the cell cycle, particularly in mitotic nuclear division and mitotic sister chromatid degradation (**Fig. 2B**). Gene set enrichment analysis (GSEA) confirmed the enrichment of the inflammatory response, highlighting the interferon-gamma and -alpha response in the H1.3 KO, while the WT condition was enriched for cell cycle-related signatures (**Fig. 2C**). Notably, compared to the H1.3 KO condition, the WT cells showed an enrichment of the DREAM complex and E2F target signatures (**Appendix Fig. S3A**). The upregulated genes in the H1.3 KO condition are involved in the interferon-gamma and -alpha response, and the downregulated genes involved in the cell cycle are highlighted in (**Appendix Fig. S3B**; **Table S6)**. To validate the functions of H1.3, we analyzed published transcriptomic data from an H1.3 KD of a human breast cancer cell line (Sancho et al., 2008) (**Table S2**). As with H1.3 KO AML cells, H1.3 KD breast cancer cells demonstrated a down-regulation of cell-cycle effectors involving E2F targets and DREAM complex signatures (**Appendix Fig. S3C**), but the interferon-gamma and alpha response signatures were not retrieved in the H1.3 KD upregulated genes. To investigate this effect further, we computed H1.3 KO inflammatory signature and tested its enrichment in the H1.4 KD and multi-H1 KD breast cancer models (Izquierdo-Bouldstridge, 2017) (**Table S2**). We showed that our H1.3 KO related “inflammatory signature” was enriched in the multi-H1 KD model (as reported in Izquierdo-Bouldstridge, 2017), suggesting a common pattern by which depletion of H1 variants influences inflammatory responses (**Appendix Fig. S3D**).

**Figure 2.**
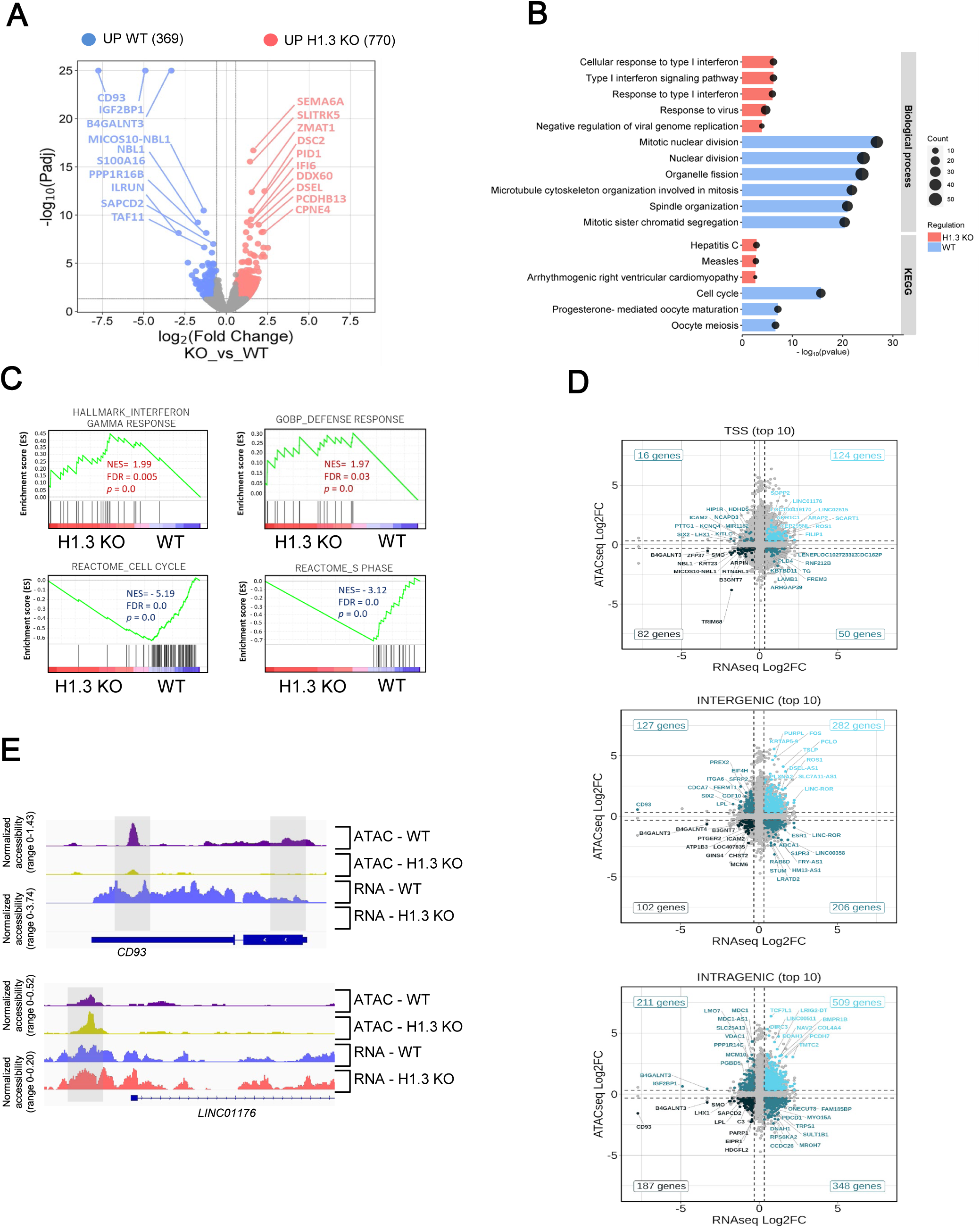
H1.3 depletion disrupts inflammation and interferon response, and cell cycle genetic programs. (**A**) Volcano plot displaying differentially expressed genes between 1 WT and 2 H1.3 KO OCI-AML3 cells with Padj < 0.05, | log2(fold change) | ≥ 0.585 (n = 3 biological replicates per cell line). Genes upregulated in the WT condition are color-coded blue, and those upregulated in the H1.3 KO condition are color-coded red. (**B**) Gene ontology (GO) enrichment and KEGG pathway analysis of upregulated genes in WT (blue) and upregulated genes in H1.3 KO (red). (**C**) Gene set enrichment analysis (GSEA) of WT and H1.3 KO OCI-AML3 cells against interferon-gamma and defense responses, cell cycle and S-phase signatures. (**D**) log2(FC) of ATAC-Seq (y-axis) plotted with the log2(FC) of RNA-Seq (x-axis) at the TSS, intragenic and intergenic genomic regions on WT and H1.3 KO OCI-AML3 cells. (**E**) Coverage plot of ATAC signals and RNA-Seq signals per condition (WT and H1.3 KO) associated with *CD93* and *LINC01176* genes. Grey boxes indicate regions with significantly different signals.

Next, to assess the potential relationship between alterations in open chromatin regions identified in ATAC-seq analysis and changes in gene expression, we crossed ATAC-seq data with our RNA-seq data. This analysis revealed four main groups of genes based on the direction of change: those with increased chromatin accessibility and upregulated expression, those with increased accessibility and downregulated expression, those with decreased accessibility and downregulated expression, and those with decreased accessibility but upregulated expression (**Fig. 2D**; **Table S7**). Among these, several genes displayed coordinated regulation of chromatin and transcription (**Table S8**). Notably, *CD93* was consistently downregulated and associated with reduced chromatin accessibility, whereas *LINC01176* was upregulated and displayed increased accessibility in the H1.3 KO condition (**Fig. 2E**).

In summary, these analyses pinpoint a role for H1.3 in the control of cell-cycle progression, aligning with observations seen in other H1 variants in breast cancer cell lines (Sancho et al., 2008). Furthermore, we revealed that the absence of H1.3 stimulates interferon signaling and inflammatory response, representing the first documented association of H1.3 histone with these pathways.

### H1.3 is associated with the H3K27me3 histone mark and its deletion remodels H1.2 localization

To further elucidate the genomic function of H1.3 in AML cells, we performed ChIP-seq to localize H1.3 in OCI-AML3 cells, using both endogenous H1.3 and ectopic V5-tagged H1.3. We first verified that the doxycycline induced V5-tagged H1.3 protein was expressed at levels similar to those of the endogenous H1.3 protein (**Appendix Fig. S4A**) and was enriched at specific chromatin regions (**Appendix Fig. S4B**). We confirmed the expected “H1 valley” around transcription start sites (TSS) for endogenous H1.3 and V5-tagged H1.3 (**Appendix Fig. S5A**), as previously reported for other H1 variants (Krishnakumar et al., 2008; Pascal et al., 2023; Salinas-Pena et al., 2024a). To further characterize the repercussion of H1.3 KO on chromatin structure, we investigated H1.3 and H1.2 genomic localization along with repressive histone marks (H3K27me3 and H3K9me3) in both WT and H1.3 KO cells (**Appendix Fig. S5B**). To measure and classify the H1 variants within the genome, we used the G-bands classification based on GC content, where GC-rich bands are linked to euchromatin and transcriptional activity, while GC-poor bands correlate with heterochromatin and repressive features (Serna-Pujol et al., 2021). Analysis of H1 distribution showed that both endogenous H1.3 and H1.3-V5-tagged were enriched in GC-rich regions, whereas H1.2 was predominantly found in GC-poor regions (**Fig. 3A**; **Appendix Fig. S5C**). This result confirms the specificity and differential nature of the genomic localization of the two histone linkers H1.2 and H1.3. Interestingly, upon H1.3 KO, the enrichment of H1.2 in GC-poor regions was reduced, suggesting a redistribution of H1.2 in the absence of H1.3 (**Fig. 3B**). To better characterize this redistribution, we analyzed the correlation between the endogenous H1.3 and H1.2 localization in WT and H1.3 KO conditions in 100-kb bins. In WT cells, H1.2 and H1.3 exhibited a modest correlation in their distribution (rs = 0.463), and H1.2 localization was mildly affected by H1.3 KO (rs=0.878), thereby confirming the specificity of action of both H1 variants (**Fig. 3C**). Noticeably, the genomic localization of H1.2 in the H1.3 KO condition exhibited a stronger correlation with H1.3 localization than in the WT condition (rs=0.748 in H1.3 KO vs rs =0.463 in WT). This observation suggested the possibility of redistribution of H1.2 to occupy sites previously occupied by H1.3 (**Fig. 3C**). Next, we compared the association of H1.2 and H1.3 with the two repressive marks, H3K9me3 and H3K27me3. As previously reported in other tissues (Kim et al., 2015; Liu et al., 2023; Salinas-Pena et al., 2024a), in AML cells, H1.2 correlated identically with both H3K27me3 and H3K9me3 (rs=0.675 and rs=0.66, respectively) (**Fig. 3D**). By contrast, H1.3 correlated poorly with the repressive H3K9me3 mark (rs=0.242) while being more correlated with H3K27me3 (rs=0.669), suggesting that H1.3 is associated preferentially with H3K27me3 along the genome (**Fig. 3D**). Interestingly, and consistent with a potential H1.2 relocalization to H1.3 sites, H1.2 gained in association with H3K27me3 (rs=0.771 in H1.3 KO vs rs=0.675 in WT) while losing in association with H3K9me3 upon H1.3 KO (rs=0.5651 in H1.3 KO vs rs=0.66 in WT) (**Fig. 3D**). A Spearman’s correlation plot is presented, summarizing our observations that H1.3 does not colocalize with H1.2 but that deleting H1.3 forced an H1.2 relocalization at H1.3-free sites (**Fig. 3E, red squares**).

**Figure 3.**
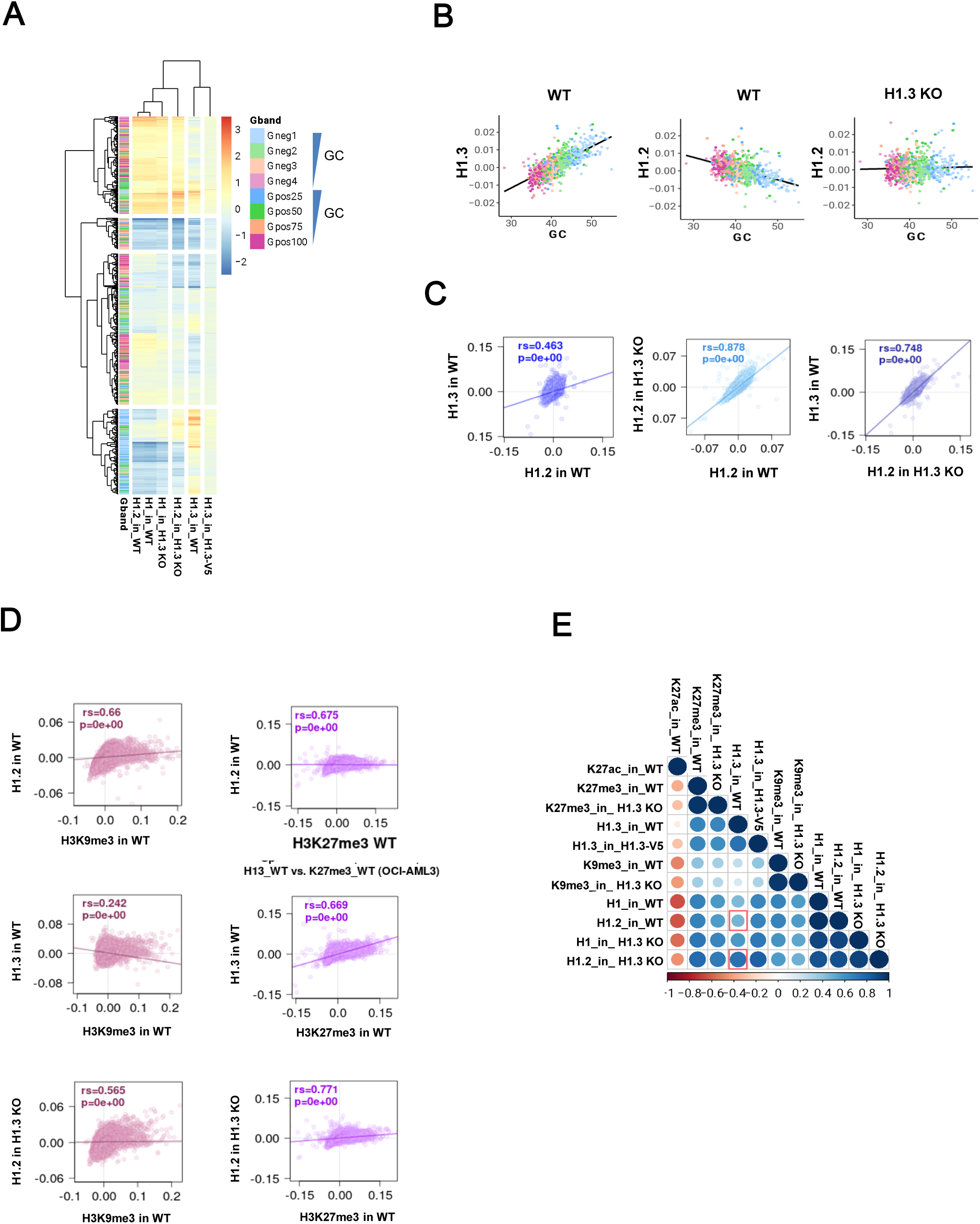
H1.3 shows distinct genomic distribution and association with repressive chromatin marks, and its loss alters H1.2 localization. (**A**) Heatmap and cluster analysis of the input-subtracted ChIP-Seq abundance (scaled) of H1 variants (OCI-AML3 cells) within Giemsa bands (G-bands). The Y-axis annotation indicates which group the G-band belongs to, as indicated in the legend. The relative GC content of the different G-bands is indicated. Data used for this analysis correspond to the averaged values obtained from two independent ChIP-seq biological replicates. (**B**) Scatter plots of the indicated H1 variant pairs (H1.2 and H1.3) input-subtracted ChIP-seq abundance within 100-kb bins of the human genome in the WT and H1.3 KO cells. The GC content at each bin is color-coded. Pearson’s correlation coefficient is shown (p < 0.001). (**C**) Spearman’s correlation plots between H1.2 and H1.3 input-subtracted ChIP-Seq signal in WT and H1.3 KO cells. (**D**) Spearman’s correlation plots between the repressive histone marks (H3K27me3 and H3K9me3) and the histone variants (H1.2 and H1.3) input-subtracted ChIP-Seq signal in WT and H1.3 KO cells. (**E**) Spearman’s correlation coefficients between all the features analyzed in 100-kb bins. Only correlations with p < 0.01 were considered (colored circles in the correlation matrices). Red squares indicate the correlation changes of H1.2 enriched bins with H1.3 enriched bins in WT and H1.3 KO cells.

Overall, these results highlight a non-random but connected genomic localization of H1.3 and H1.2, characterized by an association with different histone marks, indicating a specificity of action of histone linker variants.

### H1.3 deletion remodels H1.2 localization in regions where gene expression is deregulated

To further investigate the effect of H1.3 deletion on H1.2 localization, we focused on the top 10 % bins enriched in H1.2 signal upon H1.3 KO in comparison to WT (**Fig. 4A**). We observed a significant gain in H1.2 ChIP-seq signal in the H1.3 KO condition in the top 10% bins in comparison to random bins (**Fig. 4B**). This gain in the H1.2 signal at chromatin was also observed in more stringent bins such as the top 1%, 3% and 6.5% bins (**Appendix Fig. S6A**), supporting that H1.2 gain at chromatin is not random. The H1.3 ChIP-seq signal was markedly higher in the top bins compared with the random bins (**Fig. 4C**), suggesting that regions acquiring the H1.2 in the H1.3 KO condition were previously enriched in H1.3, but not in H1.2. This result supports the compensatory redistribution of H1.2 upon H1.3 depletion. Additionally, both H3K27me3 and H3K9me3 marks showed alterations in the top and random bins upon H1.3 KO, suggesting broader chromatin alterations beyond H1.2 redistribution (**Appendix Fig. S6B**, **C**). Next, we tested whether regions that increased H1.2 abundance upon H1.3 depletion also changed their chromatin accessibility. We observed that these bins decreased chromatin accessibility in the H1.3 KO condition; however, this decrease was not specific to the top 10 % of bins, as we also observed the same effect in the random bins (**Fig. 4D**).

**Figure 4.**
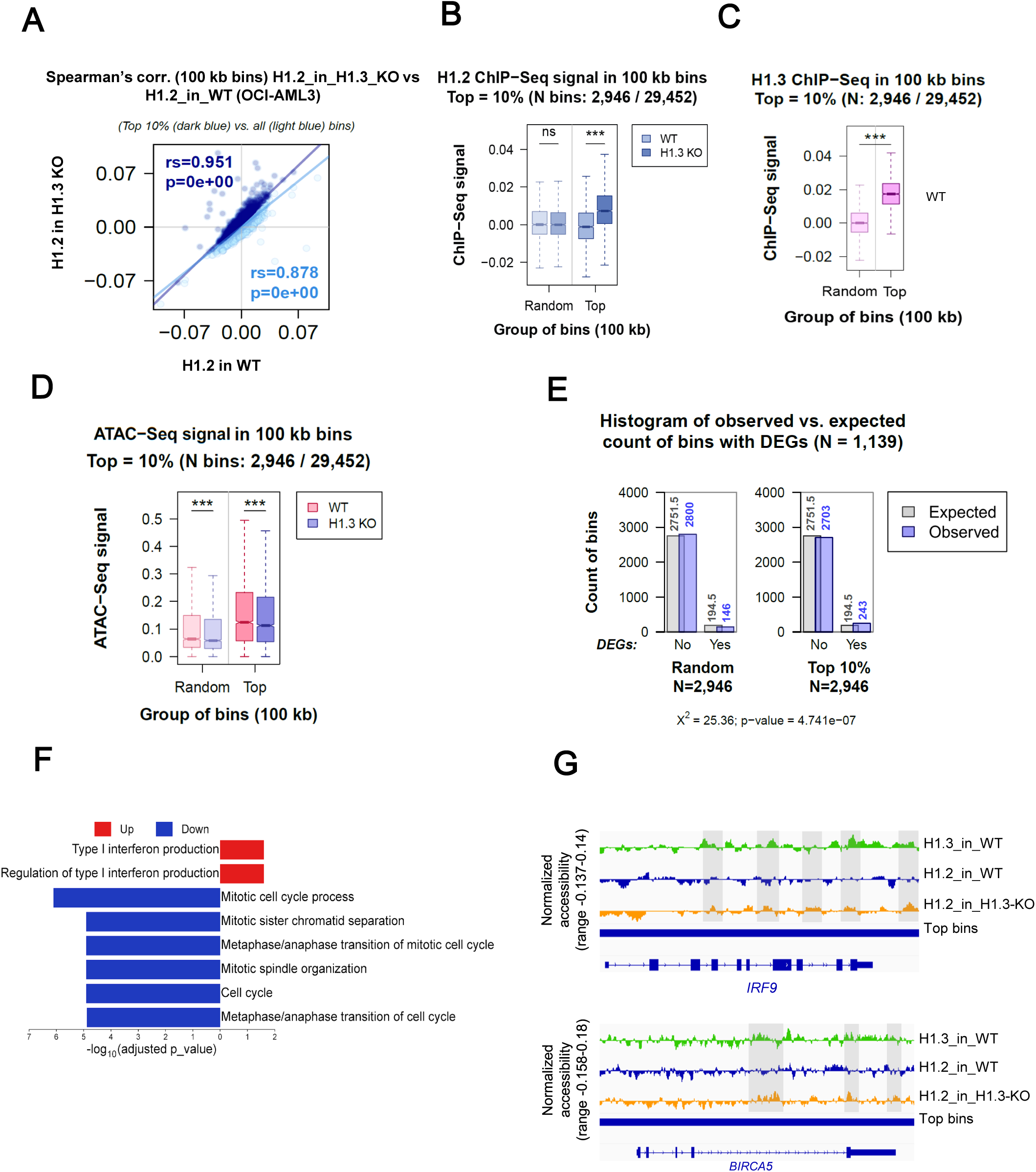
H1.3 deletion alters the genomic distribution of H1.2 in OCI-AML3 cells. (**A**) Scatter plot displaying the genome-wide changes and Spearman’s correlation of histone H1.2 ChIP-Seq signal in H1.3 KO cells compared to WT. (**B**) Boxplot showing H1.2 ChIP-seq signal in the top 10% bins where H1.2 abundance increases in the H1.3 KO condition compared to WT (labeled light blue in **A**). (**C**) Boxplot representing ChIP-seq signal for H1.3 in the top H1.2-enriched bins (from Figure 4B) compared with randomly selected bins, indicating that H1.3 previously occupied regions, gaining H1.2 in the KO. (**D**) ATAC-seq signal in top and random bins showing reduced chromatin accessibility in H1.3 KO cells. (**E**) Chi-square test of independence for testing the association of the top 10% H1.2-enriched bins with differentially expressed genes (DEGs). Out of 2946 bins, 243 overlap with DEGs, with 36 % associated with downregulated and 14 % with upregulated genes. (**F**) GO term enrichment analysis of DEGs located within the top H1.2-enriched bins. Upregulated genes are enriched for interferon signaling pathways, represented in red, while downregulated genes are associated with cell cycle regulation, represented in blue. (**G**) IGV snapshots of representative DEGs located within H1.2-enriched bins. *IRF9*, an interferon-stimulated gene upregulated in H1.3 KO cells, and *BIRC5*, a cell cycle regulator downregulated upon H1.3 depletion, are shown as examples of loci affected by H1.2 redistribution.

We then assessed whether these regions, characterized by increased H1.2 abundance and previously occupied by H1.3, were associated with changes in gene expression. We selected the top 10% of bins with the highest increase in H1.2 signal when comparing H1.3 KO versus WT conditions, as previously described, and crossed them with the DEGs. Of the 2,946 top 10% of bins, 243 (8.25% of bins) overlapped with DEGs, which was more than would be expected by chance (as 1,139 DEGs represent a 4.03% of the total number of annotated genes in the genome), indicating an association between changes in histone variant occupancy and gene expression (**Fig. 4E**). Statistical significance of this association was further confirmed by a permutation test with 10,000 iterations (**Appendix Fig. S7A**), indicating a strong association between changes in histone variant occupancy and gene expression. More precisely, 132 of these bins were associated with downregulated genes (132 genes, 35.8% of all downregulated genes), and 106 bins were associated with upregulated genes (101 genes, 13.6% of all upregulated genes). Moreover, 5 bins were associated with both upregulated and downregulated genes (**Appendix Fig. S7B**; **Table S9**). In summary, the top 10% bins enriched in H1.2 upon H1.3 KO were enriched in DEGs, especially downregulated genes, supporting the idea that H1.2 redistribution may be associated with gene repression, possibly due to its stronger repressive potential compared to H1.3. To explore the biological relevance of these changes, we performed GO enrichment analysis on DEGs located in the top H1.2-enriched bins. Notably, several of those genes were already highlighted in our global RNA-seq analysis (**Table S5**). Specifically, upregulated DEGs located within these bins were significantly enriched in interferon signaling, while downregulated DEGs were enriched in cell cycle regulation (**Fig. 4F**; **Table S10**). To provide representative examples of these chromatin changes at specific loci, we visualized ChIP-seq signals using IGV. We selected *IRF9*, an interferon-stimulated gene upregulated in the H1.3 KO condition, and *BIRC5*, a key regulator of the cell cycle that is downregulated upon H1.3 depletion (**Fig. 4G**). Additional examples of these chromatin changes at specific loci are provided in (**Appendix Fig. S8**).

Taken together, these results suggest that H1.3 depletion in AML cells promotes the redistribution of H1.2 in regions previously occupied by H1.3. Besides, this redistribution is associated with changes in gene expression, characterized by more downregulated genes than upregulated genes, and affecting pathways related to interferon signaling and cell cycle regulation.

### H1.3 depletion is associated with the upregulation of certain transposable elements (TEs) and H1.2 redistribution within these elements

H1 linker histones have been implicated in regulating transposable elements (TEs) across different systems (Choi et al., 2020; Healton et al., 2020; Salinas-Pena et al., 2024a). Here, we examined the impact of H1.3 depletion on TEs expression. Differential expression analysis revealed changes in multiple TE families, including LINEs, RC, LTR, and DNA elements, as shown in the MA plot, with several of these TEs upregulated in H1.3 KO cells (**Fig. 5A**; **Table S11**). To explore whether this effect was associated with changes in histone H1 variant distribution, we examined the abundance of H1 variants (H1.2 and H1.3) across the different TE element classes and families in WT and H1.3 KO OCI-AML3 cells using our ChIP-seq data. In WT cells, H1.3 was highly enriched at the SVA class, while H1.2 was not enriched at SVA but was more abundant at RNA elements (**Fig. 5B**). Neither variant was considerably enriched at DNA, RC, or LINE classes (**Fig. 5B**; **Appendix Fig. S9A**, **B**). Interestingly, in the absence of H1.3, the H1.2 variant showed a redistribution, with increased enrichment at SVA families in H1.3 KO cells (**Fig. 5B**, **C**). This redistribution of H1.2 in H1.3 KO cells was confirmed by meta-repeat profile analysis (**Fig. 5D**). Upon H1.3 depletion, H1.2 was also enriched at some satellite families (acro, telo, etc) where H1.3 was abundant (**Fig. 5B**, **C**). This may suggest that H1.3 was relevant at some TEs and H1.2 takes its role upon H1.3 KO. This finding may explain why the deregulation of TEs is mild and no deregulation of SVA and satellites was observed (**Fig. 5A**).

**Figure 5.**
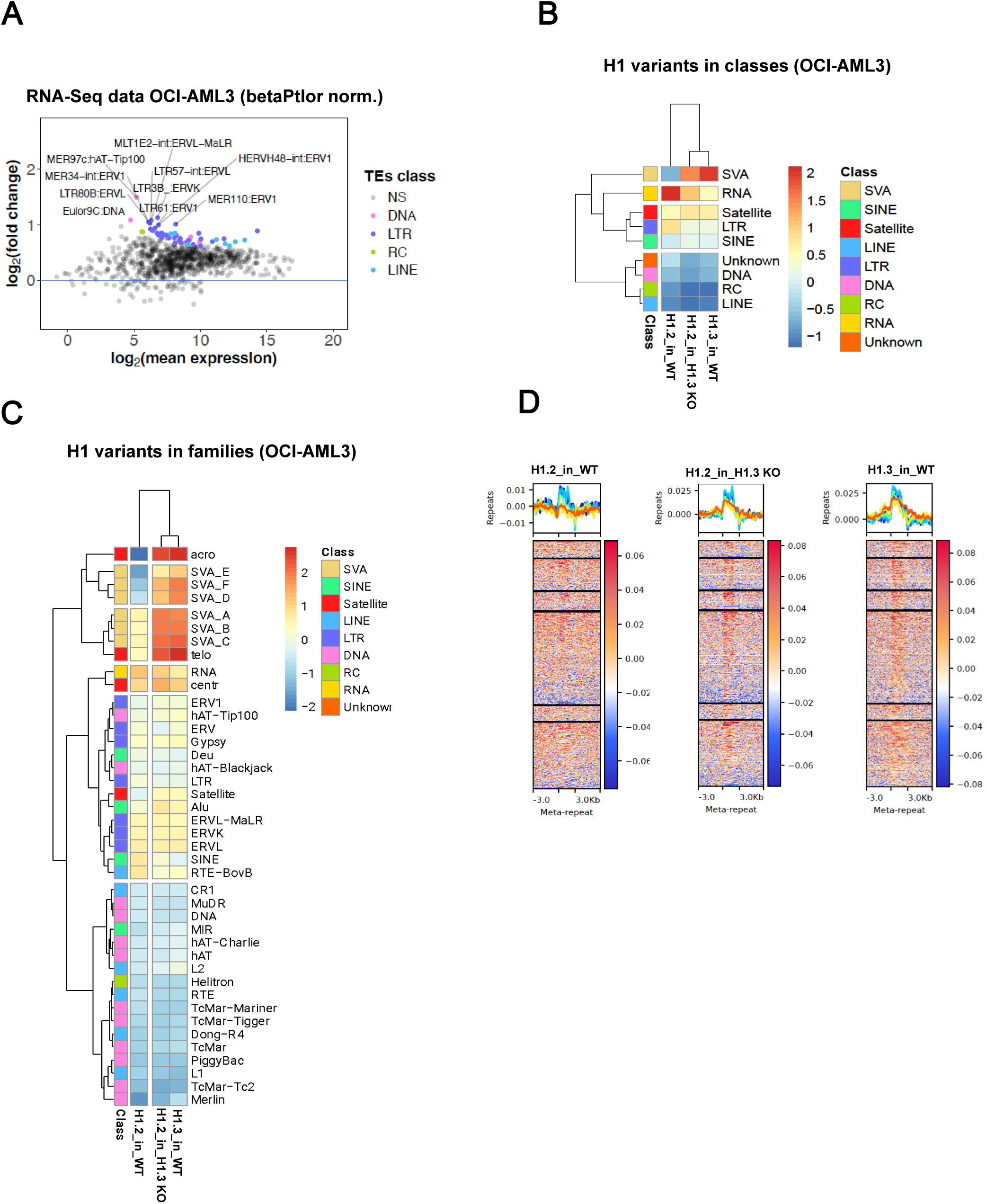
H1.3 loss is linked to transcriptional activation of transposable elements. (**A**) MA plot showing differential expression of transposable elements (TEs) in WT and H1.3 KO OCI-AML3 cells. Significantly deregulated TEs (Padj < 0.05) are highlighted and colored according to their TE class. (**B-C**) Heatmap and cluster analysis of the average input-subtracted ChIP-seq abundance of H1 variants across repeat elements families in WT and H1.3 KO cells. In **B**, each cell on the map represents the scaled average signal, with repeat families grouped by TE class (indicated on the y-axis). In **C**, Families are shown separately including the different SVA classes. (**D**) Meta-repeat profile of input-subtracted ChIP-seq signal for H1 variants at SVA repeats and their 3 kb flanking regions. All SVA copies were scaled to a uniform length for visualization. In the heatmaps, each row corresponds to a single SVA element, ordered based on the H1 binding profile in each condition.

### H1.3 KO alters G1/S transition in synchronized leukemic cells

To study the biological effect of these deregulations, we explored cell cycle regulators in more detail. First, we performed KEGG pathway analysis of cell-cycle genes, highlighting the up- and downregulated genes in H1.3 KO (**Appendix Fig. S10**) to select critical genes involved in potential G1 phase deregulation following H1.3 depletion. We confirmed the upregulation of three critical cyclin-dependent kinase inhibitors, *CDKN1A*, *CDKN2A*, and *CDKN2B*, in H1.3 KO cells (**Fig. 6A**). In addition, we observed the downregulation of *CD93* (**Fig. 6A**), a gene previously linked to leukemic stemness and cell proliferation (Iwasaki et al., 2015; Jia et al., 2022; Richards et al., 2020), in H1.3 KO cells. Next, to precise the effect of H1.3 depletion on cell cycle progression, we performed cell cycle activity tests in our cellular clones. Neither DNA content staining (Fx-violet) nor the use of the proliferation marker Ki-67 could detect a cell cycle default in the KO condition at steady state (**Appendix Fig. S11A**, **B**). We then synchronized the cells at the G1/S boundary by double-thymidine block followed by cell cycle release (Surani et al., 2021; R. C. Wang & Wang, 2022) to study how H1.3 KO alters progression through the cell cycle. As expected, T0 corresponds to cells enriched in G1 phase, T4 in S phase, and T8 in G2/M phase. By comparing the two conditions, we observed an increase in the H1.3 KO cells in G1 at the expense of the S phase compared to WT, suggesting an alteration of the G1/S transition in the absence of H1.3 (**Fig. 6B**). We did not observe any difference in the G2/M phase. The results show that the lack of H1.3 alters the progression from the G1 to the S phase of AML cells. At the molecular level, we investigated the effect of H1.3 deletion on cell cycle regulator protein expression after thymidine block. We showed that E2F1, a transcription factor driving S-phase entry, was significantly reduced in H1.3 KO cells at T0 (G1) and T4 (S), suggesting an impaired activation of the G1/S program (**Fig. 6C**). Although the RB protein level was not affected by H1.3 KO at early time points, a significant decrease of RB was detected at T8 (G2/M) in H1.3 KO cells (**Fig. 6C**). After the release of the thymidine, the phosphorylated form (pRB) was less abundant in H1.3 KO cells. Consequently, Cyclin E (CCNE1), a marker of the G1/S transition, decreased in KO cells, witnessing a default in the G1/S transition observed at 4 h after Thymidine release (T4) (**Fig. 6C**).

**Figure 6.**
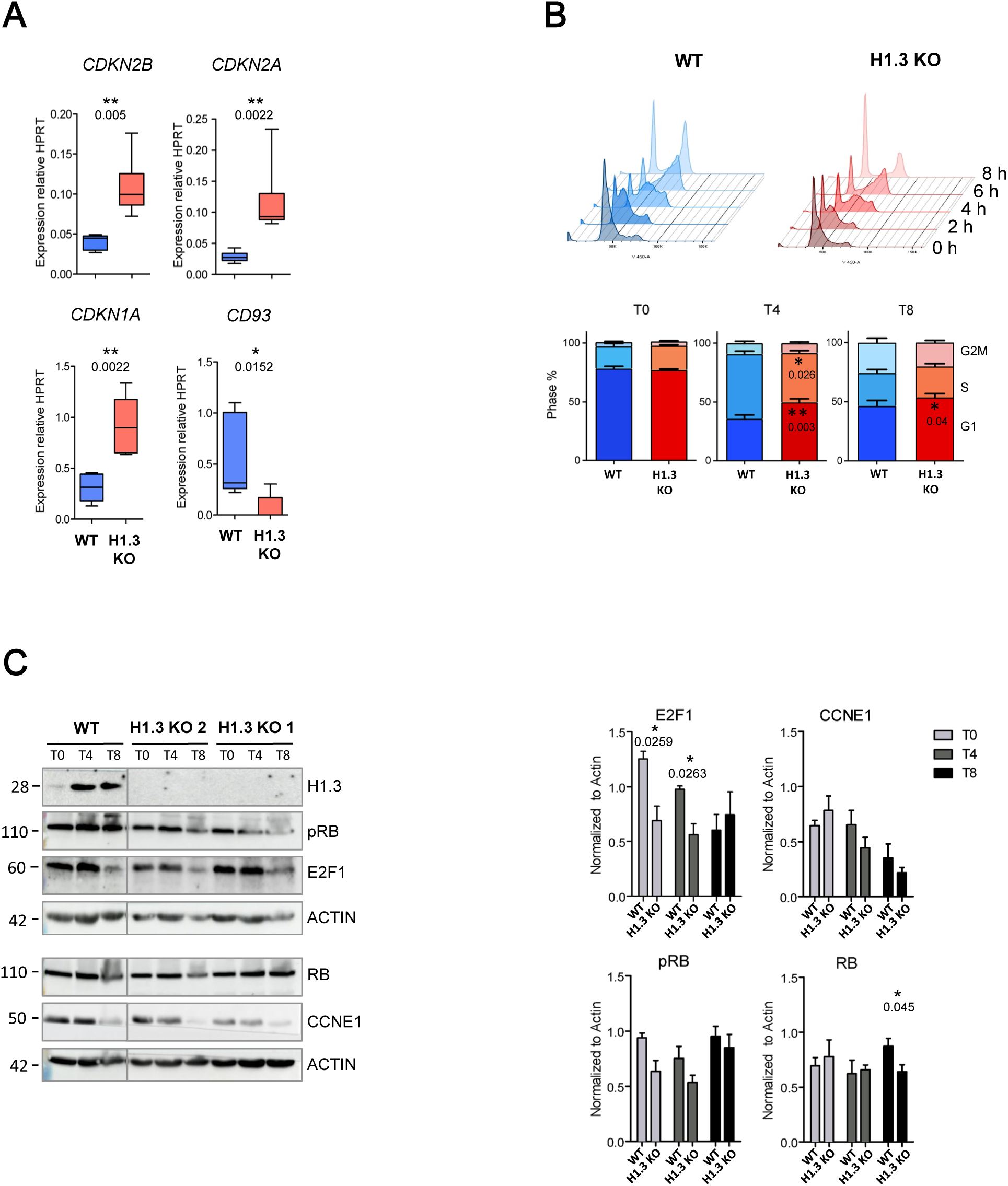
H1.3 depletion affects the G1/S transition of the cell cycle. (**A**) Analysis of cyclin inhibitor expression. Genes were quantified by RT-qPCR in 2 WT and 3 H1.3 KO independent OCI-AML3 cells. mRNA levels were normalized to the average of the housekeeping gene *HPRT*. Data represent mean ± s.e.m. of n = 6 biological replicates. Statistical significance was estimated using an unpaired two-tailed t-test * p < 0.05; **p < 0.005; ns = no significance. (**B**) Cell cycle analysis after double thymidine block release (see material and methods) of WT (blue) and H1.3 KO (red) cells. On the top panel typical Fx-Violet fluorescence profiles are shown for the two conditions at different time points (2, 4, 6 and 8). At the bottom, a comparison of the evolution of cells in the different phases of the cell cycle between WT and H1.3 KO clones at T0, T4, and T8 is shown. Data represent mean ± s.e.m. of n = 6 biological replicates. Statistical significance was estimated using an unpaired two-tailed t-test, * *p* < 0.05; **p < 0.005. (**C**) Western blot analysis of cell cycle regulators RB, phospho-RB (pRB), and Cyclin E (CCNE1) at 2, 4, and 8 hours after thymidine release in WT and H1.3 KO cells (n >= 2 biological replicates). Statistical significance was estimated using an unpaired one-tailed t-test, * *p* < 0.05.

Altogether, our analyses pointed out that the absence of H1.3 affects G1/S progression in AML cells and underlines a specific role of H1.3 in AML cells.

## DISCUSSION

The linker histone H1 family has recently gained attention across various biological processes, particularly in cancer. Our previous work, which highlighted the importance of the downregulation of the H1.3 variant in AML (Garciaz et al., 2019; Tiberi et al., 2015), led us to investigate the specific chromatin functions of H1.3 in AML. Understanding the role of H1 linker variants in chromatin remains a challenge due to the difficulty in obtaining specific antibodies for each variant compatible with ChIP-sequencing experiments. We used a recently developed specific H1.3 antibody (Salinas-Pena et al., 2024a) to characterize chromatin binding patterns of H1.3 in the OCI-AML3 cell line. In many aspects, we could demonstrate that H1.3 shares a common chromatin pattern with other H1 variants. We detected the characteristic “H1 valley” at the transcription start sites of highly expressed genes, and we witnessed an H1.3 enrichment towards the ends of gene bodies (Cao et al., 2013; Izquierdo-Bouldstridge et al., 2017; Millán-Ariño et al., 2014; Pascal et al., 2023; Serna-Pujol et al., 2022). Furthermore, H1.3 was found to be associated with the repressive H3K27me3 histone mark in a locus-specific manner. This supports its role in chromatin compaction and transcriptional regulation (Willcockson et al., 2021; Liu et al., 2023), and aligns with recent findings that linker histones are enriched in H3K27me3-marked polycomb domains (Matthews et al., 2025). However, H1.3 distribution showed some specificity, which could explain its unique role in AML cells. Notably, the chromatin distribution of H1.3 in AML cells differed from the patterns reported in breast cells (Salinas-Pena et al., 2024a), kidney cells (Pascal et al., 2023), and a mouse model (Cao et al., 2013), reflecting a context-specific contribution of H1.3 to chromatin and gene regulation. When comparing H1.3 and H1.2 distribution in the OCI-AML3 cells, we observed a weak correlation between the two variants. In addition, while H1.2 was enriched in low GC regions and showed stronger correlation with H3K9me3, a mark of constitutive heterochromatin, than H1.3 (Cao et al., 2013; Serna-Pujol et al., 2021; Bujosa et al., 2024; Kim et al., 2015), H1.3 was enriched in high GC regions, which are typically associated with open chromatin and active transcription (Serna-Pujol et al., 2021). This observation not only highlights the complexity of GC-rich regions, which can host both active genes and repressed polycomb regions (Lynch et al., 2012; Mendenhall et al., 2010) but also underscores the functional specificity of the different H1 variants.

Previous studies have shown that in a breast cancer cell line, H1.2, H1.3, and H1.5 localize to the nuclear periphery and associate with low GC regions (Salinas-Pena et al., 2024a, 2024b). Conversely, H1.4 is more uniformly distributed and enriched at high GC regions, except in cell lines that do not express H1.3 and H1.5, where H1.4 relocates to the nuclear periphery. This points to a compensatory mechanism where the chromatin localization of H1 variants adapts depending on which variants are expressed. In our AML model (OCI-AML3 cells), we noted a lower expression of H1.4 by RT-qPCR compared to the other variants. This may explain the atypical presence of H1.3 in high GC regions, suggesting that H1.3 could be fulfilling a role typically assigned to H1.4 in other cells. Moreover, in H1.3 knockout cells, H1.2 appears to redistribute from low to high GC regions, further supporting the idea of compensatory dynamics among H1 variants. Although we do not have a complete understanding of H1 variant dynamics in AML, we clearly highlighted a chromatin pattern.

In contrast to previous observations, our AML model is distinguished by the absence of compensatory upregulation of any other replication-dependent H1 variant upon loss of H1.3 (Fan et al., 2001, 2003; Luo & Brouwer, 2013). Instead, and in agreement with observations in breast cancer lines, we find an increased expression of the cell cycle-independent variant, H1.0 (Izquierdo-Bouldstridge et al., 2017; Sancho et al., 2008). Although the functional role of H1.0 in our model remains unclear, its expression could contribute to the maintenance of chromatin integrity in the absence of H1.3. Another notable compensatory effect is the partial redistribution of H1.2 in H1.3 KO cells, particularly at sites freed by H1.3, which could result in different transcriptional outcomes. H1.2 partially replaces H1.3 at certain loci, specifically those associated with the repressive H3K27me3 mark. This redistribution moves H1.2 away from its usual domains with low GC content and high H3K9me3, suggesting a chromatin reorganization favouring greater accessibility. Interestingly, the regions where H1.2 accumulates frequently correspond to loci whose expression is modified, particularly genes that are repressed in the H1.3 KO.

All these observations support the hypothesis that there is a direct link between deregulation of chromatin binding of H1 variants and transcription. They also support that H1 variants are not equivalent in terms of gene regulation; their impact depends on their association with chromatin features (H3K27me3, H3K9me3, and GC content) and their interaction with regulatory complexes. The main transcriptional changes observed upon H1.3 deletion concern interferon signaling pathways, the cell cycle, and the regulation of transposable elements (TEs). Chromatin activation of the immune response linked to H1 disruption has been described in other systems, often via derepression of satellites and TEs and accumulation of cytoplasmic double-stranded RNAs (Izquierdo-Bouldstridge et al., 2017; Salinas-Pena et al., 2024a; Vlasova et al., 2025). In our case, H1.2 is recruited to TE loci previously occupied by H1.3, and specific families of TEs are activated upon loss of H1.3. However, this activation is not universal and, for example, LINEs and SINEs elements remain unaffected. These observations suggest that H1.3 contributes to the repression of a distinct subset of TEs that H1.2 cannot fully compensate for.

The altered distribution of H1.2 could also contribute to the cell cycle defects observed in H1.3-deficient cells by directly disrupting regulatory chromatin domains or their interactions. This is confirmed by the repression of cell cycle-related genes and the overexpression of CDK inhibitors (*CDKN1A*, *CDKN2A*, and *CDKN2B*), as well as by the decreased of CCNE1, E2F1 protein and RB phosphorylation levels, blocking G1/S progression. These results align with the findings of Sancho et al., who observed transcriptional downregulation of cell cycle genes by microarray analysis following H1.3 deregulation (Sancho et al., 2008). In addition, H1.2 has been proposed as a direct regulator of the cycle through an interaction with the RB protein (Lai et al., 2021; Munro et al., 2017). Linked to the chromatin impact of H1.3 KO and its transcriptional signature, cell cycle progression was impacted in the absence of H1.3. We observed an accumulation of cells in G0/G1 following synchronization, indicative of a dysregulation of the G1/S transition, particularly the repression of targets of the DREAM and E2F complexes (Bujarrabal-Dueso et al., 2023; Kent & Leone, 2019; Sadasivam & DeCaprio, 2013). We have also identified strong repression of CD93, involved in leukemic stem cell proliferation (Iwasaki et al., 2015; Jia et al., 2022), with reduced chromatin accessibility at its locus, suggesting direct control by H1.3. Finally, the downregulation of MCM complex components suggests impairment of replication initiation or replicative stress, as observed with the depletion of other H1 variants (Almeida et al., 2018; Fernández-Justel et al., 2022; Ozgencil et al., 2023). A similar chromatin-based increase in replicative stress was recently reported upon H1 depletion (Ingham et al., 2025), where chromatin decompaction enhanced topoisomerase II, supporting a key role for linker histones in maintaining chromatin architecture that supports accurate DNA replication and S-phase entry (Ingham et al., 2025). It is essential to note that our studies were conducted in *NPM1*-mutated cells; thus, the influence of *NPM1*-mut in our results cannot be excluded. The association between NPM1 and H1.3 remains unclear, emphasizing this as a focal point for our future studies to better understand *NPM1* mutation, the linker histone H1.3, and the overall AML patient H3K27me3*HIST1*-positive phenotype.

In summary, we provide new insights into the mechanisms of action and chromatin specificities of the histone linker variant H1.3. We highlight its involvement in cell cycle regulation and interferon signaling in AML cells. While H1.3 has been described as an oncogene in pancreatic cancer (Bauden et al., 2017) and as a tumor suppressor in ovarian cancer (Medrzycki et al., 2014), our data indicate that H1.3 supports leukemic cell proliferation through epigenetic cell cycle regulation. This work supports the idea that H1 variants display tissue-specific roles in cancer. It provides a basis for future studies to explore the complex interplay between H1.3, chromatin organization, and gene regulation in the context of AML to highlight its potential as a therapeutic target in AML where it is dysregulated.

## MATERIAL AND METHODS

### Cell lines and culture conditions

Human cell lines (OCI-AML3 and derived clones) were grown in MEMα medium supplemented with 10% fetal bovine serum, 100 U/mL penicillin, and 100 U/mL streptomycin under standard culture conditions (37°C, 5% CO2). Cell lines were regularly verified to be Mycoplasma-free using PCR detection and thawed to perform experiments within a 6-to-10-passage period.

**For CRISPR-Cas9-mediated H1.3 KO**, we created a customized editing construct using an H1.3 guide RNA (gRNA-AAGAAGGCAGGCGCAACTGC), which was cloned into the px458 CRISPR-Cas9 expression plasmid (Addgene #48138). The plasmid (px458-H1.3-gRNA) was transfected into the target cells by electroporation using the Amaxa Kit V (program X-001) according to the manufacturer’s protocol. Transfected cells were identified based on the presence of green fluorescent protein (GFP) and cloned by sorting one cell per well on 96-well plates using the BD FACS Aria III cell sorter. The loss of H1.3 was evaluated in the different clones by Western blot. Mutational profiles of the clones were obtained by Sanger sequencing of the region targeted by the gRNA (Eurofins Genomics).

**For OCI-AML3 cells expressing a V5-tagged H1.3**, we cloned the H1.3 DNA into a doxycycline-inducible plasmid pLix-403 (Addgene #41395) using the GATEWAY strategy and transfected the pLix403-H1.3-V5 construct into OCI-AML3 cells by electroporation using the Amaxa Kit V (program X-001). Transfected cells were selected 48 h after electroporation with puromycin 2 µg/mL. Expression of the H1.3-V5 was induced with 1 µg/mL doxycycline and assessed by Western blot.

Doxycycline (Sigma; #D5207) was resuspended at 10 mg/ml in DMSO and stored at −20°C. Intermediate dilutions were prepared in a culture medium before being added to the cells.

### Protein extraction and western blot

Cells were washed and resuspended with ice-cold PBS and lysed in 2x Laemmli buffer (62.5 mM Tris-HCl, pH 6.8, 2% SDS, 10% glycerol, 5% β-mercaptoethanol, and 0.5% bromophenol blue) at 95°C for 8 minutes. To extract histone proteins from DNA, lysates were sonicated using an ultrasonic homogenizer (Bioruptor^TM^ Next Gen; Diagenode; 5 cycles, 30 seconds of sonication followed by 30 seconds of rest) and the soluble fraction was collected by centrifugation at 16.100g for 15 minutes at 4°C. The samples were loaded and separated under denaturing conditions using SDS-PAGE with the Mini-Protean® Tetra System, Bio-Rad, at 100 V with electrophoresis buffer (25 mM Tris pH 8.3, 192 mM glycine, and 0.1% SDS). After separation, proteins were transferred to nitrocellulose membranes using a transfer buffer (25 mM Tris, pH 8.3, 192 mM glycine, and 10% ethanol) at 200 mA for 2 hours. The membranes were blocked with either 5% milk or 1% BSA in TBS-T (Tween at 0.05%) for 1 hour at room temperature (RT). Subsequently, the membranes were incubated with different primary antibodies overnight at 4°C, followed by three 10-minute washes with TBS-T at RT. Then, the membranes were incubated with a secondary antibody (mouse or rabbit, 1/10000) for 1 hour at RT and were revealed with chemiluminescent reagent (ECL 1:1)-ECL Prime™ or ECL Select™ (GE Healthcare) according to the detected protein, for 5 minutes and then developed using the Amersham Imager 680.

The primary antibodies used were: H1.0 (#56695 Santa Cruz Biotechnology), H1.2 (#4086 Abcam), H1.3 (#203948 Abcam), H1.4 (##41328S Cell signaling), H1.5 (#18208 Abcam), H1 (#39708 Active Motif), pRB (#81805 Cell Signaling), RB (#93095 Cell Signaling), E2F1 (#3742S Cell Signaling); Cyclin E (#HE12 Sc 247 Santa Cruz); V5-tag (Abcam; #15828), actin (#A1978 Sigma) and tubulin (#T8535 Sigma). The polyclonal goat anti-mouse or anti-rabbit immunoglobulins HRP were used as secondary antibodies (Dako; P0447, P0448, respectively). Band quantification from the obtained images was performed using the ImageJ Lab software.

### RNA extraction and RT-qPCR

Total RNA was extracted using the RNeasy® Mini Kit from Qiagen (74104). RNA elution was carried out using RNase-free water and RNA concentration was measured at 260 nm using a spectrophotometer (Nanodrop 2000, ThermoFisher Scientific). The A260/A280 ratio was used to assess RNA purity, with pure RNA having an A260/A280 ratio within the range of 1.9 to 2.1.

Total RNA was reverse-transcribed using the Transcription High Fidelity cDNA Kit (Roche, #5091284001), following the manufacturer’s protocol. For the polymerase chain reaction, SYBR Green Fast Master Mix from Bio-Rad was used, along with a set of specific gene primers (**Supplementary Table S1**). The mRNA expression was normalized to the housekeeping gene HPRT. Relative expression levels were determined using the ΔCT (Cycle threshold) method, where ΔCT = Ct gene - Ct HPRT, and the relative gene expression was calculated as 2^(−ΔCT).

### Cell cycle analysis

#### FxCycle™ Violet Stain

Cells were seeded at a density of 1×10^6^ cells per mL in a 24-well plate, 24 hours before the analysis. Cells were collected, fixed with cold 70% ethanol and stained with FxCycle™ Violet Stain (Invitrogen™ #F10347). The cells were washed and resuspended in a FACS buffer solution and cell cycle analysis was performed using a BD-LSRII cytometer (BD Biosciences), allowing us to analyze the different phases of the cell cycle based on DNA content.

#### Ki67 proliferation assay

Cells were seeded at a density of 1×10^6^ cells per mL in a 24-well plate. 24 hours before the analysis, cells were collected and fixed by adding 100 µL of BD Cytofix/Cytoperm^TM^ (BD Biosciences #554722) containing formaldehyde, followed by incubation in the dark for 20 minutes at RT. Cells were collected and resuspended in 100 µL of BD Cytoperm Permeabilization Buffer Plus (BD Biosciences, #561651) for 10 min at RT. The permeabilized cells were thoroughly resuspended in 100 µL of 1× BD Perm/Wash buffer (BD Biosciences #554723) before the Ki67 intracellular staining. A titrated anti-Ki67 fluorochrome-conjugated antibody was added and incubated at 4°C overnight, and cell cycle analysis was performed using a BD-LSRII cytometer.

#### Cell cycle synchronization by double thymidine blocking

Cells were synchronized in the late G1/early S phase using a double thymidine block method. To initiate the synchronization process, 300.000 cells/mL were seeded in T-100 dishes four hours prior to treatment. The first thymidine block was started by the addition of thymidine (Sigma; #T9250-5G, 100 mM stock solution stored at −20°C) to the dishes at a final concentration of 2 mM for 12 hours. Cells were then washed 3 times with PBS and cultured in normal cell maintenance media for 12 hours, followed by a second 12-hour thymidine (2 mM) block. Cells were released from the double thymidine block and collected every 2 hours and processed for flow cytometry.

### RNA Sequencing (RNA-seq)

#### Data generation

Total RNA was extracted from the WT OCI-AML3 parental cell line and two H1.3 KOs (KO clone #1 and KO clone #2) using the RNeasy® Mini Kit (Qiagen; 74104), as previously described. Each experimental condition includes three biological replicates. Libraries were prepared with rRNA depletion with a Kapa mRNA HyperPrep kit (Roche, #KR1352-v7.21) and sequenced using the Illumina NovaSeq6000 platform, 100-bp paired-end reads, at 50 million reads per sample at Genomics and Bioinformatics Platform Marseille (Marseille, France).

### Data analysis

#### Alignment

Sequencing quality control was determined using the FastQC tool (http://www.bioinformatics.babraham.ac.uk/projects/fastqc/) and aggregated across samples using MultiQC (v1.7) (Ewels et al., 2016). Reads with a Phred quality score of less than 30 were filtered out and mapped to the human (Hg38) reference genome using Subread-align (v1.5.0) (Liao et al., 2013) with default parameters. Gene expression quantification was performed by counting mapped tags at the gene level using featureCounts (Liao et al., 2014).

#### Differential analysis

Differentially expressed genes were identified using DESeq2 (v1.26.0) (Love et al., 2014) and were selected according to the following criteria: Padj <=0.05, absolute (FC) >1.5. For IGV visualization, mapped reads were converted into Bigwig using the Deep Tools suite (v2.2.4) (*bamCoverage --scaleFactor library_size_factor --binSize 5*) (Ramírez et al., 2016).

#### Functional enrichment analysis

The Gene Ontology (GO) analysis was performed using clusterProfiler with a Padj <= 0.05 and by considering a background. Then, the genes were split into two groups according to their fold-change (UP genes: (FC) >= 1.5, DOWN genes: (FC) <= 1.5). The GO categories chosen were Biological Process (BP) and KEGG pathways.

The gene set enrichment analysis (GSEA) was performed with different gene lists as input to the GSEA software (v4.3.2). For the H1.3 KO gene lists, we considered the differentially expressed genes between H1.3 KO and WT conditions with Padj <= 0.05 and (FC) => 1.5) and ranked them according to their (FC) of all genes detected in the RNA-seq experiments, which were used for the ranked gene lists.

For the H1.3 knock-down (KD) and multi-H1 KD gene lists, we considered the H1.3 KD DEGs obtained from GSE12299 Microarray (Sancho et al. 2008) and the dataset from GSE83277 RNA-seq (Izquierdo-Bouldstridge et al. 2017) (**Supplementary Table S2**), respectively, with Pvalue <= 0.05 and (FC) >1.4 and ranked them according to their (FC).

For the H1.3 KO specific inflammatory signature, we considered H1.3 KO DEG (Pvalue <= 0.05 and (FC) => 1.4) common to the GOBP immune response, inflammatory response and defense response gene lists. Volcano plots, GO enrichment, and Cnetplot figures were plotted by https://www.bioinformatics.com.cn/en, a free online platform for data analysis and visualization. KEGG pathways were created with Pathway-based data integration and visualization (https://pathview.uncc.edu/) (W. Luo et al., 2017; W. Luo & Brouwer, 2013). Genes ID were obtained from https://www.genetolist.com.

### ChIP Sequencing (ChIP-seq)

#### Data generation

We used protocols as previously described in (Koubi et al., 2018). In brief, 5×10^6^ OCI-AML3 cells (from WT and H1.3 KO clones) and 20×10^6^ H1.3-V5 OCI-AML3 cells were fixed with 1% formaldehyde for 8 min. After successive washes, cells were lysed with Lysis Buffer 1 (50 mM HEPES, 1 mM EDTA, 140 mM NaCl, 0.75% NP40, 10 % Glycerol, 0.25% Triton X100), and Lysis Buffer 2 (200 mM NaCl, 10 mM Tris-HCl pH8, 1 mM EDTA, 0.5 mM EGTA) and chromatin extracted in Sonication Buffer (0.5% Laurylsarcosine, 10 mM Tris-HCl pH8, 1mM EDTA, 0.5 mM EGTA, 100 mM NaCl, 0.1 % NaDoc). After sonication using the Bioruptor® Pico (Diagenode) to obtain DNA fragments with an average size of 300 bp, chromatin was immunoprecipitated overnight at 4°C with magnetic beads pre-incubated with the following antibodies: H3K27ac (Active Motif; #39133), H3K27me3 (Cell Signaling; #9733S), H3K9me3 (Active Motif; #39161), H1.3 (Abcam; #203948), H1.2 (Abcam; #4086), H1 (Active Motif; #39707), V5-tag (Abcam; #15828) and rabbit-IgG (Cell signaling; #2729S), used as a control. Immunoprecipitations were washed with the following combinations of wash buffers: W1 (1% Triton X100, 0.1% NaDOC, 150 mM NaCl, 10 mM Tris-HCL pH8), W2 (0.5% NP40, 0.5% Triton X100, 0.5% NaDOC, 150 mM NaCl, 10 mM Tris-HCL pH8), W3 (0.7% Triton X100, 0.1% NaDOC, 250 mM NaCl, 10 mM Tris-HCL pH8), W4 (0.5% NP40, 0.5% NaDOC, 250 mM LiCl, 20mM Tris-HCL pH8, 1mM EDTA, W5 (0.1% NP40, 150 mM NaCl, 20 mM Tris-HCL pH8, 1 mM EDTA), W6 (20 mM Tris-HCL pH8, 1 mM EDTA) and DNA was purified with an I-Pure V2 Kit (Diagenode).

The primers used for verification by ChIP-qPCR are listed in **Supplementary Table S3**.

Libraries were generated using the MicroPlex Library Preparation Kit (Diagenode) following the manufacturer’s instructions and the quality and size distribution of library fragments were analyzed on a Bioanalyzer 2100 system (Agilent). Sequencing was performed using the Illumina Nextseq500, 75-pb paired-end reads with 30 millions reads per sample at the Theories and Approaches of Genomic Complexity platform (Marseille, France) (for H1.3-V5 and H3K27ac ChIP) and using the Illumina Novaseq6000 platform, 100-bp paired-end reads, at 50 million reads per sample at Genomics and Bioinformatics Platform Marseille (Marseille, France) for WT and H1.3 KO clones.

### Data analysis

#### Alignment

Sequencing quality control was determined using the FastQC tool (http://www.bioinformatics.babraham.ac.uk/projects/fastqc/). Paired-end reads were trimmed using Trimmomatic (Bolger et al., 2014) and aligned to the human hg19 reference genome using Bowtie2 (v2.3.5.1) (Langmead & Salzberg, 2012) with default options. Unmapped reads, duplicates, and low-quality alignments were filtered out using SAMtools (v1.9) (Li et al., 2009) with the flag 3844. **BedGraph and BigWigs**. The resulting BAM files were sorted, and input-subtracted, counts per million (CPM)-normalized signal tracks were generated in bedGraph and bigWig formats using deepTools (Ramírez et al., 2016) (bamCompare --operation subtract --normalizeUsing CPM -- scaleFactorsMethod None). BEDTools (Quinlan & Hall, 2010) was employed to quantify the average ChIP-Seq signal of the samples within specific genomic regions of interest (i.e., G-bands, TEs, and 100-kb bins).

#### Downstream analysis

Heatmaps of histone H1 variants were generated using the R package *pheatmap*, with Euclidean distance and complete cluster method used for clustering both rows and columns. Scatter plots were produced using base R, and Spearman’s correlation was used to assess associations between histone H1 variants and PTMs. Box plots were also created using base R, and the Wilcoxon signed-rank test was applied to compare ChIP-seq signals between WT and H1.3 KO conditions within the same subset of regions. Correlation matrices were computed using the *cor()* function in R (method = “spearman”) and visualized with the *corrplot* package. ChIP-seq profiles of H1 variants at meta-repeats were constructed and visualized using deepTools (v3.5.1) (Ramírez et al., 2016) (commands: computeMatrix, plotHeatmap, plotProfile). All regions were scaled to the same length using the scale-regions mode.

#### Genome annotation and segmentations

To evaluate H1 variants abundance, different chromatin segmentations were used. Eight groups of Giemsa bands (G-bands) were defined according to (Serna-Pujol et al., 2021). Briefly, G-bands were classified as G positive (Gpos25 to Gpos100, according to its intensity upon Giemsa staining), and G-negative (unstained), which were further divided into four groups according to their GC content (Gneg1 to Gneg4, from high to low GC content).

#### TE meta-repeat profiles and heatmaps

For repetitive elements analyses, the TE transcripts (v2.1.4) annotation was used (Jin et al., 2015). Annotation includes nine different classes of repeats, including: LINE, SINE, LTR, DNA, Satellite, Other (SVA), Unknown, RC (Rolling-Circle), and RNA. Repeats with unsure classification named with a ‘?’ at the end of the family or class name (e.g. SINE?) were excluded from the analysis. Repeats overlapping problematic regions defined in ENCODE BlackList (Amemiya et al., 2019) were also excluded from the analysis. For analysis performed at the TE classes, family and group level (i.e., meta-repeat profiles and heatmaps), multi-mapping reads were used as recently incorporated TE copies have not diverged enough to efficiently assign a unique position.

### ATAC Sequencing (ATAC-seq)

#### Data generation

ATAC-seq was performed with the Diagenode ATAC-seq kit; (#C01080001) following the manufacturer’s instructions. In brief, 5×10^5^ OCI-AML3 cells from WT and H1.3 KO (clones #1 and #2) were harvested and resuspended in 500 µL of cold PBS containing protease inhibitors. Nuclei extraction was performed by adding to the cell pellet 50 µL of cold ATAC Lysis Buffer 1 containing 2% Digitonin for 90 seconds. Nuclei were collected by centrifugation, and a transposition reaction was performed by adding 50 µL of transposition mixture (containing Tagmentation buffer, Tagmentase, 2% Digitonin, 10% Tween20, PBS and Nuclease-free water) for 30 min at 37°C. DNA was isolated using supplied spin columns and eluted in 12 µL of DNA elution buffer. Transposed DNA fragments were amplified using 8 UDI for Tagmented libraries (Diagenode; #C01011035) and libraries were purified using AMPure XP beads (Beckman Coulter; #A63881).

Libraries were generated using the MicroPlex Library Preparation Kit (Diagenode) following the manufacturer’s instructions and the quality and size distribution of library fragments were analyzed on a Bioanalyzer 2100 system (Agilent). Sequencing was performed using the Illumina Novaseq6000 platform, 100 bp paired-end reads, at Genomics and Bioinformatics Platform Marseille (Marseille, France).

### Data analysis

#### Alignment

Sequencing quality control was determined using the FastQC tool (http://www.bioinformatics.babraham.ac.uk/projects/fastqc/) and aggregated across samples using MultiQC (v1.7) (Ewels et al., 2016). Reads with a Phred quality score of less than 30 were filtered out and mapped to the human (Hg38) reference genome using default parameters of Bowtie2 (v2.3.4.3) (Langmead et al., 2012). Only mapped reads were filtered using samtools (v1.9) (Li et al., 2009) (*view; -exclude-flags 4*). Duplicate tags were removed using Picard Tools (v2.20.2) (https://broadinstitute.github.io/picard/) (*MarkDuplicates; REMOVE_DUPLICATES = true*), and mapped tags were processed for further analysis.

#### BigWigs

Mapped reads were normalized by their library size and converted into bigWigs using the deepTools suite (v3.2.1) (Ramírez et al., 2016) (*bamCoverage --scaleFactor library_size_factor --binSize 5*). Reads were extended using values previously computed by deepTools *(bamPEFragmentSize*) and used through the --*extendReads option*. BigWig files from multiple replicates were merged into a single file using a custom bash script (*bigWigMerge*). All resulting files were visualized using the Integrative Genomics Viewer (IGV; v2.5.2) (Robinson et al., 2011).

#### Peak calling and differential analysis

High confidence binding sites were determined using MACS2 peak caller (v2.1.2) (Zhang et al., 2008) in broad mode (*broad-cutoff=0.05 -q 0.01 -- nomodel*). Each sample’s fragment size was previously computed by deepTools (*bamPEFragmentSize*) and used through the --*extsize option*. Coordinates shared by at least 2 samples (i.e. coverage >= 2) were selectively retained and merged into unified coordinates (*bedtools merge*). Mapped tags were then counted within these consensus coordinates using featureCounts (v1.6.4) (Liao et al., 2014) and normalized using DESeq2 (*estimateSizeFactors*) (v1.26.0) (Love et al., 2014). Finally, differential analysis was performed (*DESeq, minReplicatesForReplace = Inf, betaPrior = T)*.

#### Genomic distribution of annotated peaks

Genomic distribution was performed by Galaxy software (http://usegalaxy.org/) using the ChIPseeker tool. Significant peak changes were filtered with a P value < 0.05 and (FC) > 1. We used the nearest gene research to identify the genes that would be impacted by the difference in accessibility.

#### Gene Ontology analysis

Gene Ontology (GO) analysis was performed using clusterProfiler with a Padj < 0.05 and by considering a background. Enriched GO terms were determined using only the genes that exhibited significant modulation (Pvalue <= 0.05). GO figures were done using https://www.bioinformatics.com.cn/en a free online platform for data analysis and visualization.

### Multiomics analysis

#### RNA-seq and ATAC-seq

Differential analyses data from OCI-AML3 bulk RNA-seq and ATAC-seq were merged using a custom script. Briefly, DESeq2 output peaks from ATAC-seq differential analysis were annotated using both Homer (v4.11) (Heinz et al., 2010) (*annotatePeaks.pl*) and ChIPseeker (v1.34.0) (Wang et al., 2022) (*annotatePeak*) annotation tools. Only TSS peaks were retained for further analysis. Bulk RNA-seq and ATAC-seq data were then merged regarding their gene names. ATAC-seq and RNA-seq log2 fold-changes were plotted against each other using ggplot2 (v3.4.1). Genes were highlighted using filtering thresholds regarding their relative p-values and log2 fold-changes. Only the top 10 candidates per quadrant were labelled using ggrepel (v0.9.3) (*geom_text_repel*).

#### ChIP-seq, RNA-seq and ATAC-seq

The relationship between data from OCI-AML3 bulk RNA-seq, ATAC-seq, and ChIP-seq within 100-kb bins was assessed using a custom R script. First, bins were ranked by the log₂ fold-change in H1.2 ChIP-seq signal (KO vs. WT), and the top 10% (N = 2,946) were selected. To assess DEG enrichment in H1.2-associated regions, the 1,139 identified DEGs were mapped to 100-kb genomic bins using BEDTools. We quantified the number of bins containing at least one DEG in both the top bins and a subset of randomly selected bins (N = 2,946), and the association between H1.2-enriched bins and the presence of DEGs was evaluated using bar plots and a chi-square test. Statistical significance was further confirmed by a permutation test with 10,000 random reshufflings of bin labels. To assess ATAC-seq enrichment within 100-kb bins, the average genome-wide ATAC-seq signal in both WT and H1.3 KO conditions was mapped to the 100-kb bins using BEDTools. We then compared the average ATAC-seq abundance in top versus random bins using box plots and evaluated the statistical significance of the differences between WT and H1.3 KO conditions with a Wilcoxon signed-rank test.

### Statistical analysis

Statistical analyses were done using R software (version 2.15.2) (The Comprehensive R Archive Network. http://www.cran.r-project.org/) and Prism 9 (Graph Pad Software) and the significance of the differences between groups was determined by unpaired t-Test, Mann–Whitney test or exact Fisher test. Data was presented as the median ± SEM.

## DATA AVAILABILITY SECTION

ATAC-seq datasets are deposited in the GEO database under accession number GSE302140, CHIP-seq datasets are deposited in the GEO database under accession number GSE302143, RNA-seq datasets are deposited in the GEO database under accession number GSE302144.

## ACKNOWLEDGMENTS

The authors thank the Flow Cytometry and Bioinformatics platforms of the Centre de Recherche en Cancérologie de Marseille for their technical support. The Genomics and Bioinformatics Platform of La Timone and the Theories and Approaches of Genomic Complexity platform (Marseille, France) are acknowledged for RNA-seq library preparation and sequencing of RNA, ChIP, and ATAC samples. The authors also thank Christophe Lachaud and the members of the Duprez laboratory for their valuable discussions. Albert Jordan is acknowledged for providing the endogenous H1.3 antibody. The pDNOR-zeo plasmid was a gift from F. Lembo.

## FUDINGS

This work was supported by the Ligue Nationale Contre le Cancer; the Institut National de la Santé et de la Recherche Médicale (INSERM); the Centre National de la Recherche Scientifique (CNRS); the Canceropôle Provence Alpes Côte d’Azur, the Institute for Cancer and Immunology (Aix-Marseille University), by a grant from l’Institut National du Cancer (PRT-K16-071 to E.D.), which also funded LNG; a European Union’s Horizon 2020 Research and Innovation Program (Marie Skłodowska-Curie grant # 813091 ARCH, age-related changes in hematopoiesis (that funded CT), by the Spanish Ministry of Science, Innovation and Universities and FEDER, EU (grant number PID2023-146239OB-I00/AEI/10.13039/501100011033).

We also acknowledge the Generalitat de Catalunya Suport Grups de Recerca AGAUR (grant number 2017-SGR-597).

## AUTHOR CONTRIBUTIONS

**Clara Tellez-Quijorna:** Investigation; Methodology; Formal analysis; Visualization; Writing— original draft; Writing—review and editing. **Lia N’Guyen:** Investigation; Methodology; Formal analysis; Visualization; Writing—review and editing. **Núria Serna-Pujol:** Investigation; Methodology; Formal analysis; Visualization; Writing—review and editing. **Maeva Rodies:** Investigation; Validation, Visualization. **Julien Vernerey:** Data curation, Formal analysis; Visualization. **Albert Jordan:** Methodology; Formal analysis; Supervision; Funding acquisition; Writing—review and editing. **Estelle Duprez:** Conceptualization; Supervision; Funding acquisition; Investigation; Methodology; Project administration; Writing—review and editing.

