## Supplemental figures for "H1.3 depletion in AML cells prompts H1.2 redistribution, chromatin remodeling and cell cycle defects"

### Appendix Figure S1

## A

| H1.3 KO | Allele 1 | Effect | Allele 2 | Effect |
|---|---|---|---|---|
| Clone 1 | Del 84-99 (16) | Frameshift | Del 84-99 (16) | Frameshift |
| Clone 2 | Del 90 (1) | Frameshift | Del 90-92 (2) | Frameshift |

## B

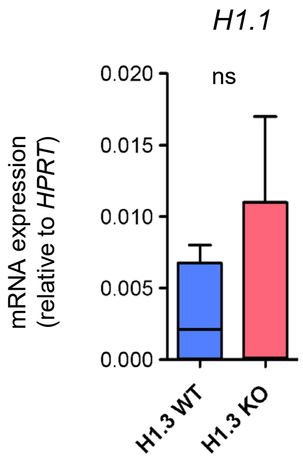


**Figure S1. Characterization of H1.3 KO OCI-AML3 cells. (A)** Table presenting the deletions generated after CRISPR-Cas9. **(B)** Gene expression of *H1.1* in OCI-AML3 (WT) and H1.3 KO clones. mRNA levels were normalized to the average of the housekeeping gene *HPRT*. Experiment represents 2 WTs and 2 KO with each sample being tested in triplicate. Statistical significance was estimated using an unpaired two-tailed t-test; ns = no significative.

### Appendix Figure S2

A

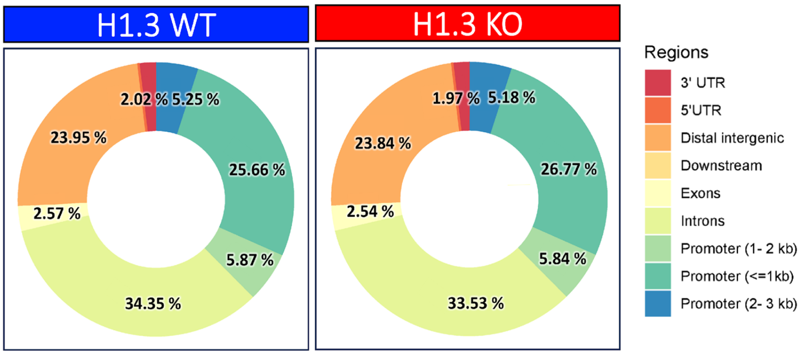


B

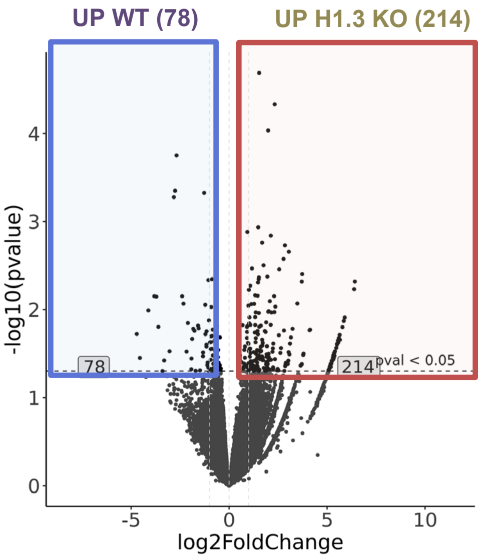


**Figure S2. The impact on chromatin accessibility in the H1.3 KO OCI-AML3 cells. (A)** Distribution in the different genomic regions of ATAC-seq peaks obtained in WT and H1.3 KO cells. **(B)** Volcano plot of differential peaks ($p < 0.05$; $|\log2(FC)| \geq 0.585$). The upregulated peaks in WT are underlined in blue, and the upregulated in H1.3 KO condition are underlined in red.

### Appendix Figure S3

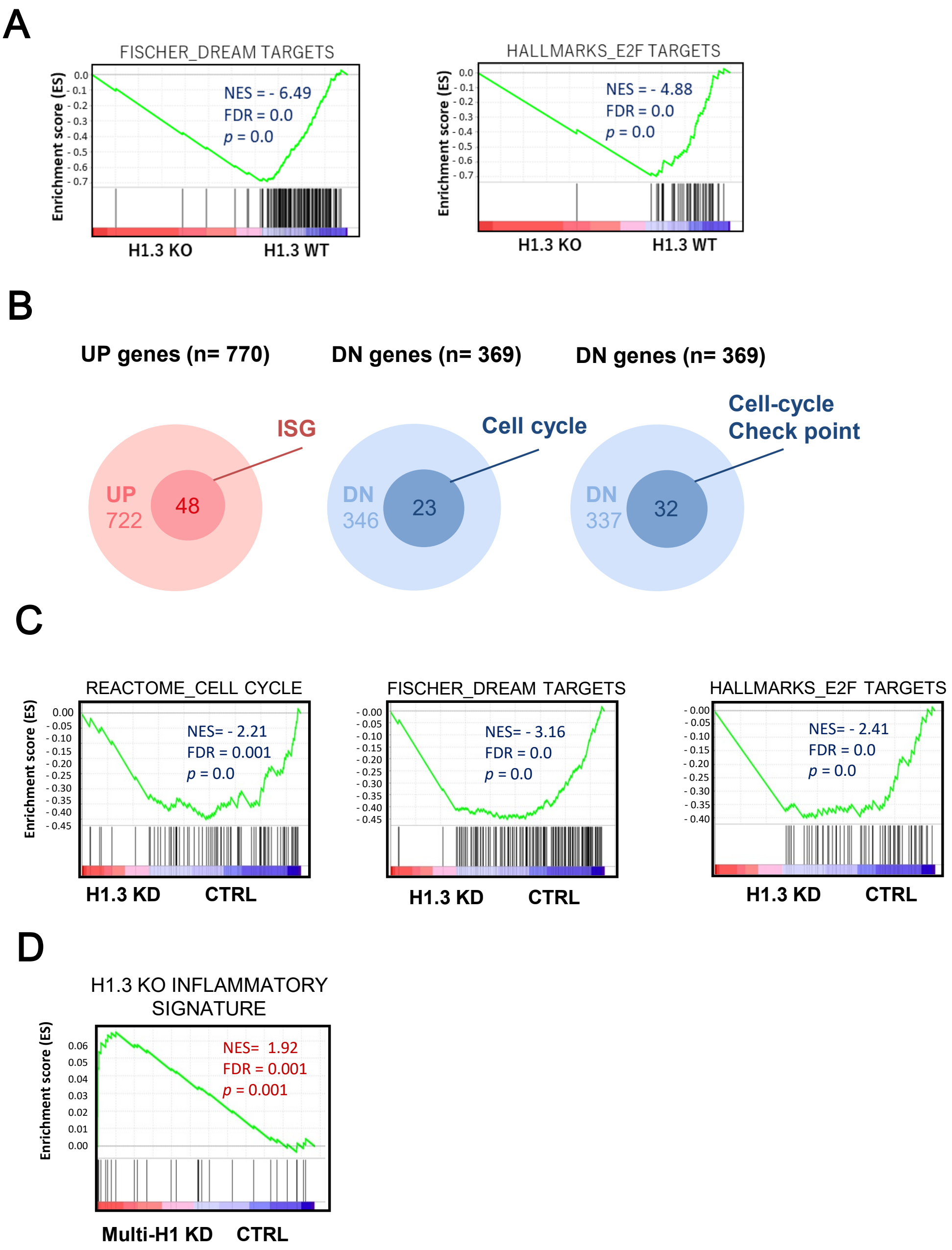


**Figure S3. H1.3 depletion or downregulation impacts cell cycle and inflammatory gene signatures. (A)** GSEA of WT and H1.3 KO cells against DREAM complex and E2F targets. **(B)** Venn diagrams illustrating the overlap between differentially expressed genes in H1.3 KO cells and functionally annotated gene sets. Left: overlap between upregulated genes (UP) and interferon-stimulated genes (ISGs), highlighting ISGs upregulated in the KO condition. Middle: overlap between downregulated genes (DN) and KEGG cell cycle genes, showing cell cycle-related genes downregulated in KO cells. Right: overlap between downregulated genes (DN) and genes involved in cell cycle checkpoints, highlighting checkpoint-related genes repressed upon H1.3 loss. **(C)** GSEA of the H1.3 knock-down induced with dox (H1.3 KD) and empty vector (CTRL) human breast cancer cells (Sancho et al., 2008) against Cell cycle, DREAM complex and E2F target signatures. **(D)** GSEA of the multi-H1 KD and empty vector (CTRL) human breast cancer cells (Izquierdo et al. 2017) against our specific H1.3 KO Inflammatory signature.

### Appendix Figure S4

A

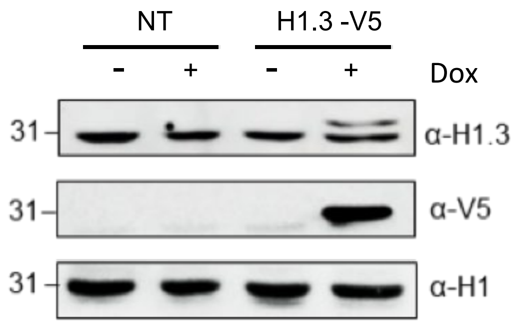


B

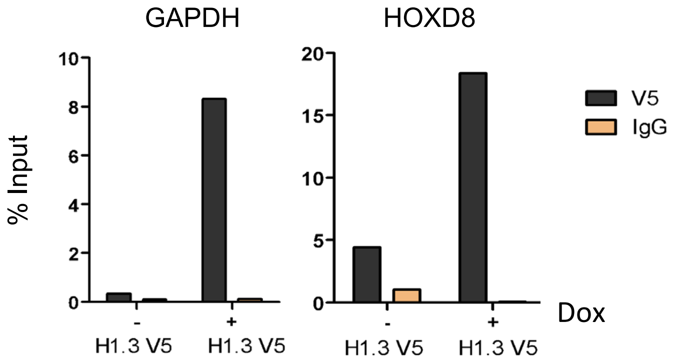


**Figure S4. Characterisation of V5-tagged H1.3-expressing OCI-AML3 cells. (A)** Western blot showing the exogenous H1.3-V5 protein in OCI-AML3 cells (H1.3-V5) following doxycycline induction (Dox, 1 µg/ml). Untransfected OCI-AML3 cells (NT) were used as a control. (**B**) V5 ChIP-qPCR experiment in OCI-AML3 H1.3-V5 cells, either induced or not with doxycycline (Dox ; 1 µg/ml).

#### Appendix Figure S5

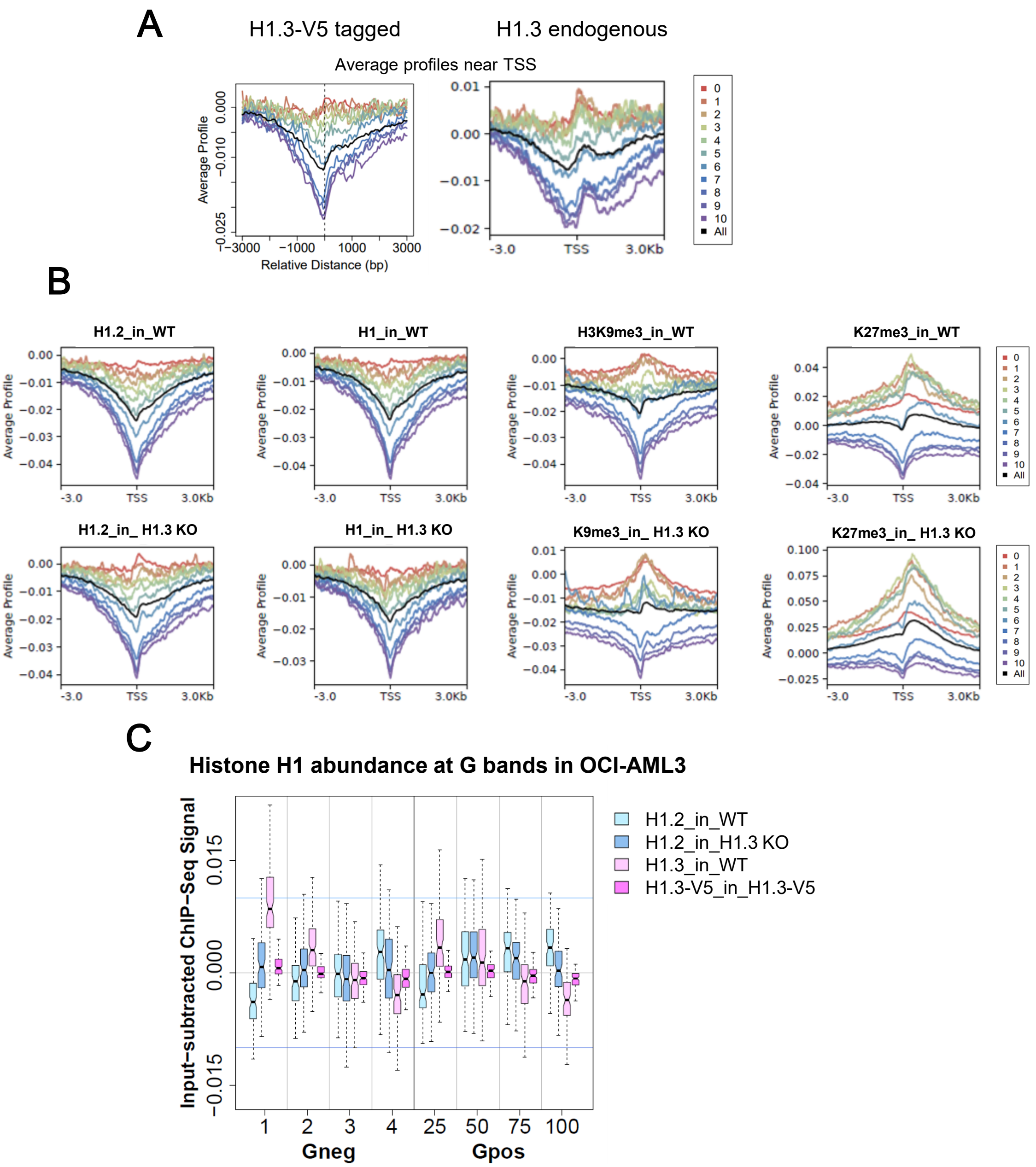


**Figure S5. ChIP-Seq comparison of H1.3-V5 tagged and endogenous H1.3 chromatin distribution. (A)** H1 variants input-subtracted ChIP-Seq average profile around gene transcriptional start site (TSS). Expressed genes are divided into 10 equal groups, each containing 10% of the total expressed genes, according to their basal gene expression on RNA-Seq experiments. Group 0 includes nonexpressed genes. Average H1 profile for all genes is shown in black. **(B)** The average input-subtracted of H1.2, H1 and histone marks ChIP-Seq regarding the relative distance to TSS for all transcripts classified according to expression in 10 groups containing the same number of transcripts, from 0-non expressed to 10-highly expressed. **(C)** Box plots showing the average input-subtracted ChIP-Seq signal of H1.3-V5 tagged, the endogenous H1.3 and H1.2 at each group of G-bands, for each band type in WT and H1.3 KO OCI-AML3 cells.

### Appendix Figure S6

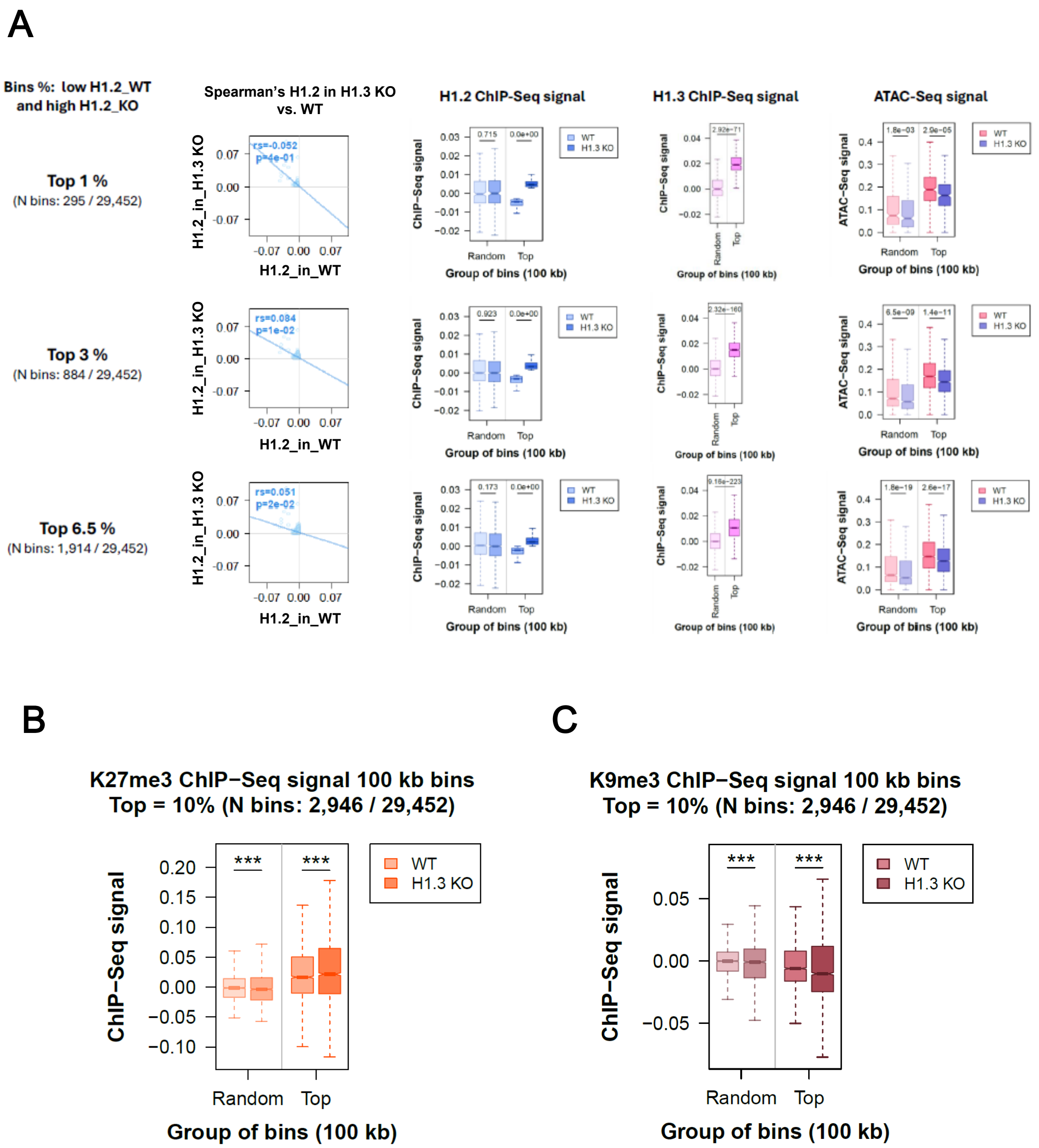


**Figure S6. Characterization of genomic regions gaining H1.2 upon H1.3 depletion. (A)** Plots showing H1.2 signal enrichment across additional top-ranked bins beyond the top 10% (top 1%, top 3% and top 6.5 %), defined by increased H1.2 levels in H1.3 KO compared to WT cells. **(B)** Average ChIP-seq signal profiles for the repressive histone mark H3K27me3 across top and random bins. **(C)** Average ChIP-seq signal profiles for H3K9me3 across the same top and random bins.

### Appendix Figure S7

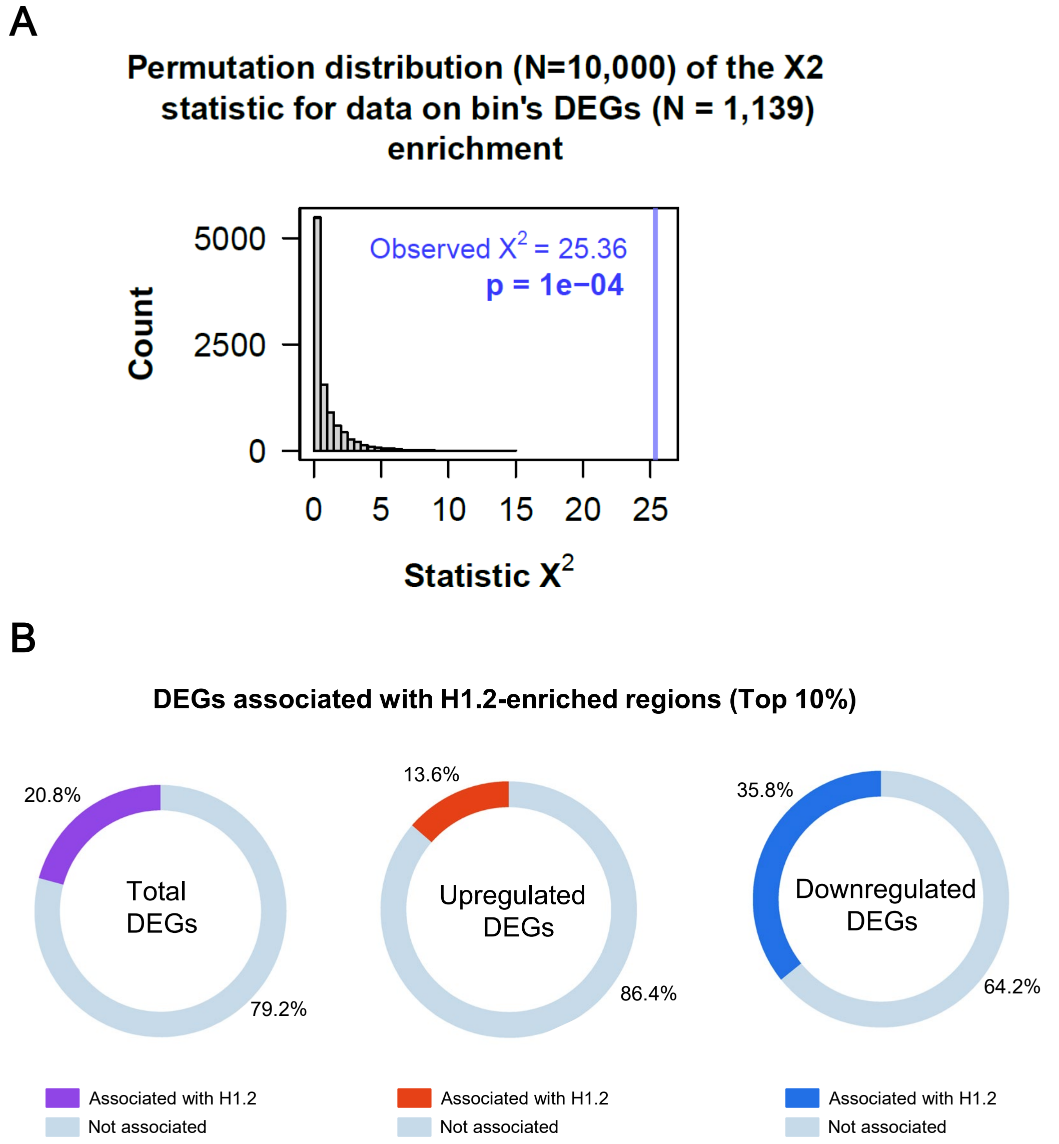


**Figure S7. Association between H1.2-enriched regions and differentially expressed genes (DEGs). (A)** Histogram showing the distribution of the $\chi^2$ statistic from the permutation test assessing DEG enrichment in top versus random bins. A total of 10,000 permutations were performed. The blue vertical bar indicates the observed $\chi^2$ value obtained from the chi-square test using the actual counts of DEGs within top and random bins. **(B)** Donut charts illustrating the proportion of DEGs overlapping with H1.2-enriched regions. Left panel: from the total DEGs (n = 1139), 243 genes (20.8%, in purple) overlapped with H1.2-enriched bins. Middle panel: among the upregulated DEGs (n = 770), 105 genes (13.6%, in red) were associated with H1.2-enriched regions. Right panel: among the downregulated DEGs (n = 369), 132 genes (35.8%, in blue) overlapped with H1.2-enriched regions.

### Appendix Figure S8

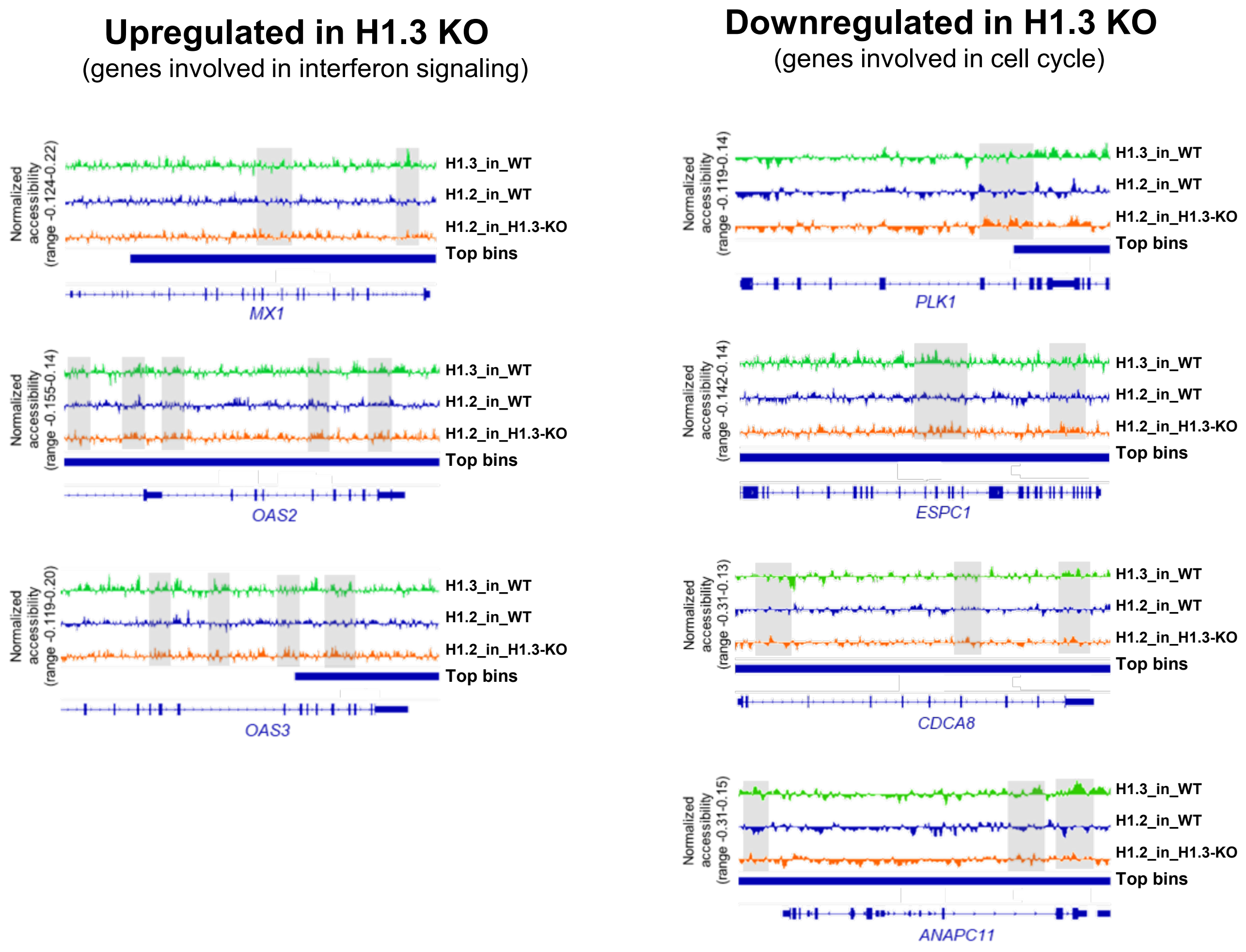


**Figure S8. Representative IGV tracks showing H1.2 redistribution at genes involved in interferon signaling and cell cycle regulation.** Screenshots from IGV displaying ChIP-seq signal tracks for H1.2 in WT and H1.3 KO cells at representative gene loci. On the left, loci correspond to interferon-stimulated genes, and on the right, to cell cycle–related genes. These genes were previously identified as differentially expressed in our RNA-seq analysis (Supplementary Figure S3B and Supplementary Table S6) and are also found within the top 10% of genomic bins showing increased H1.2 occupancy in the KO condition (Supplementary Table S9).

### Appendix Figure S9

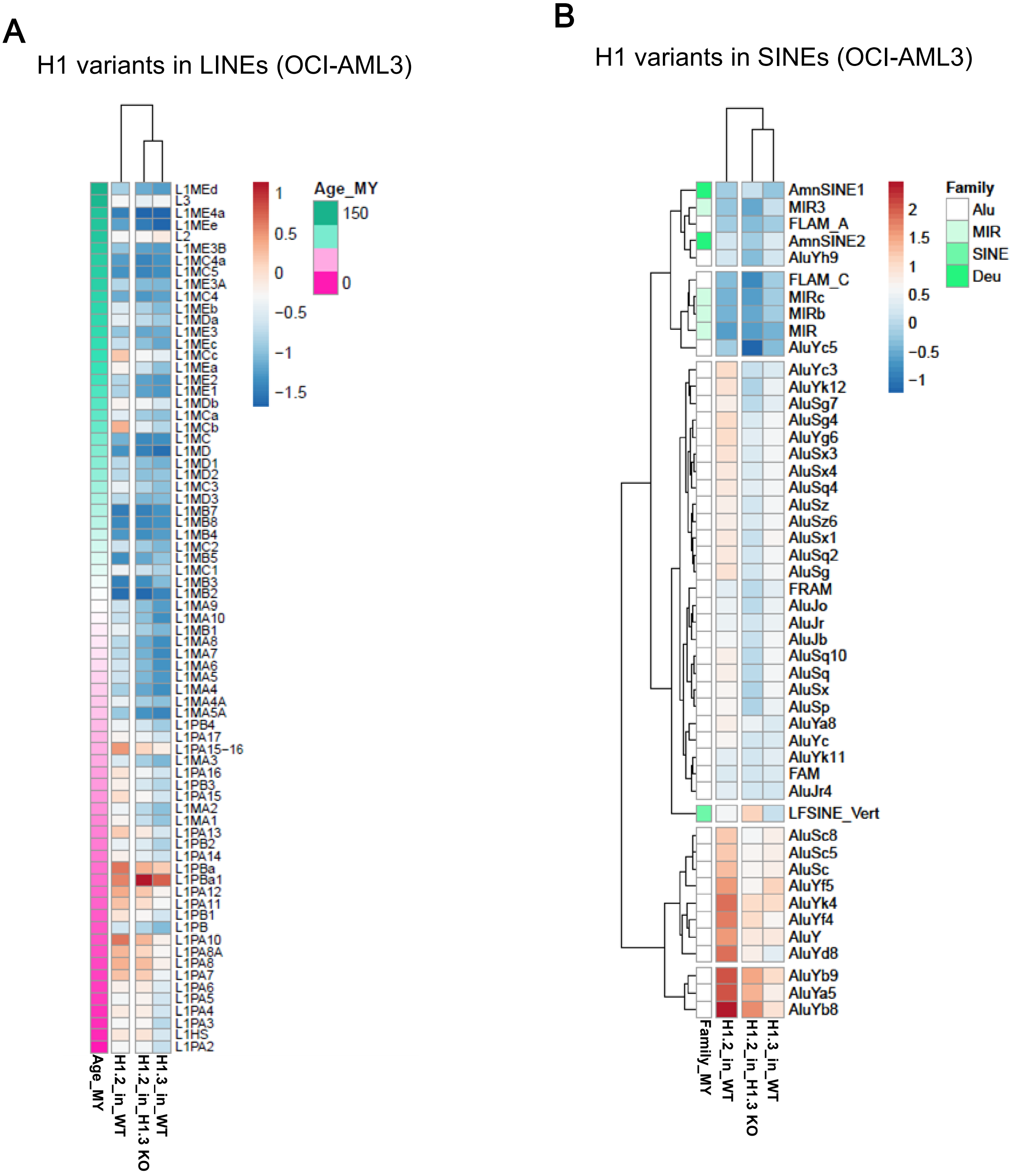


**Figure S9. Distribution of H1 Variants at LINEs and SINEs Repeats. (A)** Heatmap and cluster analysis of the average input-subtracted ChIP-Seq abundance (scaled) of H1 variants within selected groups of LINE elements (N = 73) ordered chronologically. **(B)** Heatmap and cluster analysis of the average input-subtracted ChIP-Seq abundance (scaled) of H1 variants within the groups of SINE repeats (N = 48) belonging to the four families.

### Appendix Figure S10

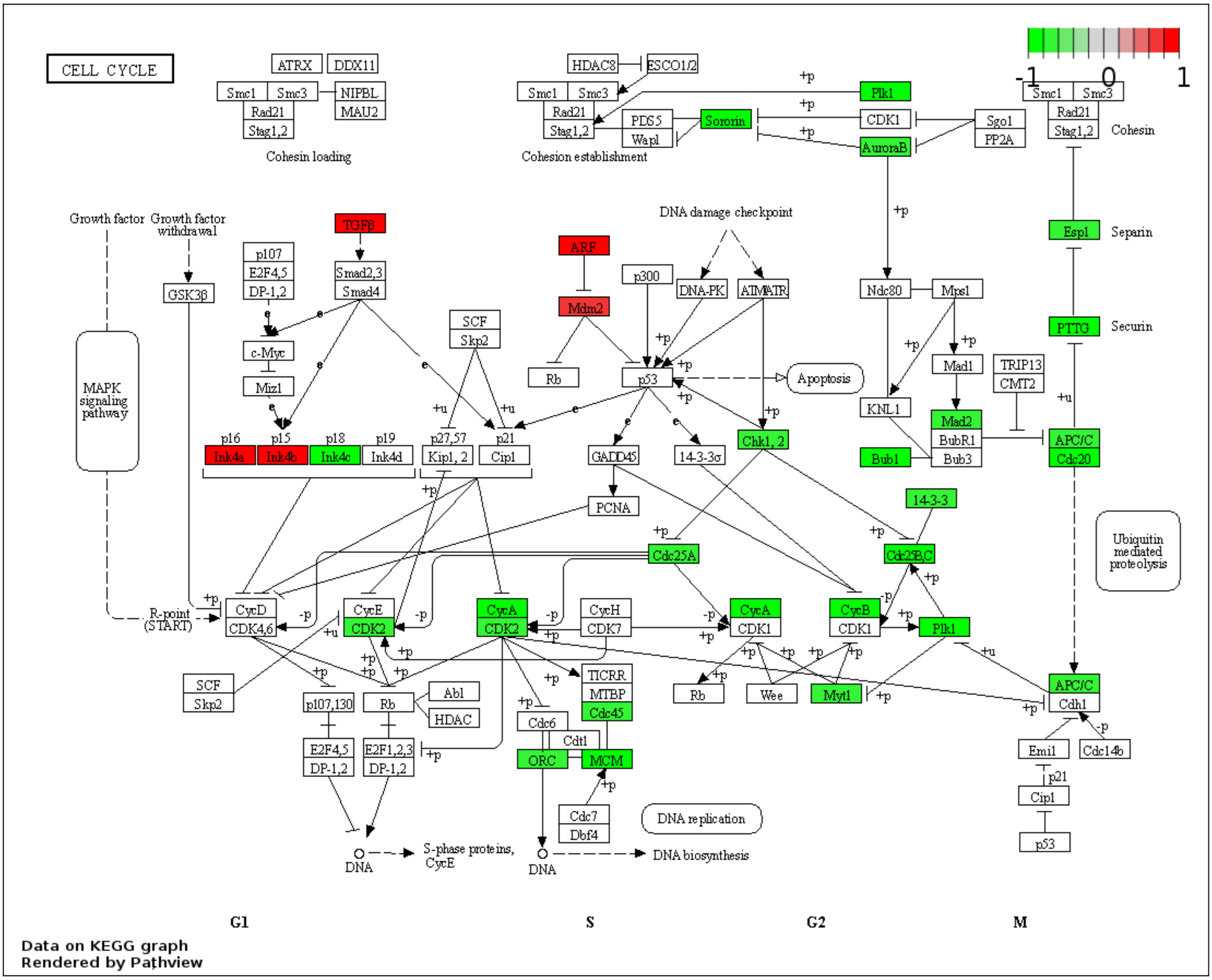

**Figure S10. Pathview visualization of differentially expressed genes of KEGG Cell Cycle pathway.** Pathview of DEG from RNA seq analysis on Cell cycle KEGG graph. The genes up regulated in the WT condition are highlighted in green, the genes down regulated in the H1.3 KO condition are highlighted in red. KEGG pathways were created with Pathway-based data integration and visualization (https://pathview.uncc.edu/) (W. Luo et al., 2017; W. Luo & Brouwer, 2013).

### Appendix Figure S11

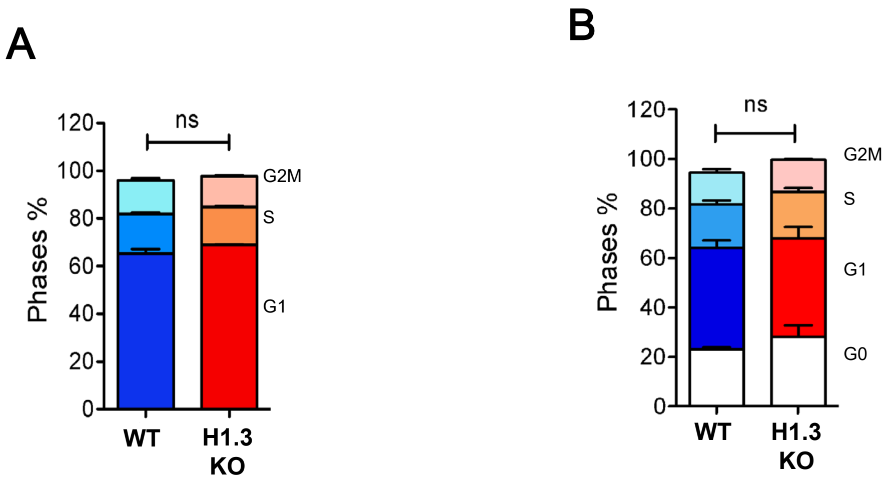


**Figure S11. Cell Cycle phase distribution in H1.3 WT and KO cells analyzed by flow cytometry.** Cell cycle analysis of WT (blue) and H1.3 KO (red) cells. Histograms of cell cycle phases identified by flow cytometry with (A) Fx Violet staining and (B) Ki67 and Fx Violet staining (n = 3 biological replicates). Values are expressed in ± SD. Statistical significance was estimated using an unpaired two-tailed t-test, ns: non-significant.
